# Spike-in–normalised single-cell RNA-seq reveals cell-type-specific transcriptional repression during ageing

**DOI:** 10.64898/2026.06.25.733584

**Authors:** Idálio de Jesus Viegas, Cyril Lagger, João Pedro de Magalhães

## Abstract

Transcriptome analyses are widely used for biomarker discovery and to gain insights into normal processes and diseases. Age-related changes in gene expression inferred from RNA-seq are typically reported relative to the transcriptome composition using library-size normalisation. As such, absolute changes in transcript abundance with age remain poorly characterised. Here, using external spike-in normalisation in the Tabula Muris Senis dataset, we quantify age-related variation in total mRNA content and gene expression across mouse cell types. We observe widespread changes in total mRNA abundance, with decreases predominantly in non-immune cell types and increases predominantly in immune cell types. In parallel, the number of genes expressed declines across most cell types, including immune populations. Differential expression analysis based on spike-in–normalised counts identifies genes consistently downregulated across cell types, enriched for functions in RNA metabolism and protein processing. Furthermore, genes downregulated during ageing and during proliferation arrest show partial overlap, suggesting that these transcriptional changes may share regulatory processes. Together, these results are consistent with a general repression of transcriptional and metabolic activity with age, modulated by immune-specific responses.

**Highlights:**

- Total mRNA content decreases with age in several non-immune cell types and increases in most immune cell types.
- The number of detected genes decreases with age in most cell types.
- Genes downregulated based on absolute normalisation are associated with RNA and protein metabolism.
- Ageing-associated transcriptional repression overlaps with gene expression programs linked to proliferation arrest.

## Introduction

Ageing is a progressive loss of biological function that increases frailty and leads to an exponential increase in mortality. The complexity and gradual nature of ageing challenge our ability to understand and design effective longevity interventions[1]. A variety of experimental and computational techniques are currently being leveraged to shed new light on this complex phenomenon[2]. Among them, transcriptomic methods enable simultaneous measurement of gene expression across thousands of genes. An early meta-analysis of age-related gene expression profiles from bulk microarray studies identified upregulation of genes associated with immune response, inflammation, and lysosomal function, and downregulation of genes related to mitochondrial function, cell cycle, and apoptosis[3]. Following these initial findings, transcriptomics has been further applied to numerous ageing studies, including bulk RNA sequencing (RNA-seq), single-cell RNA sequencing (scRNA-seq), and spatial transcriptomics (Spatial-seq)[4].

Transcriptomic studies typically focus on identifying differentially expressed genes based on relative expression levels, while exploring global mRNA abundance and the overall number of detected genes remains less common. Indeed, the total mRNA content of each sample is typically biased by experimental noise arising from capture and sequencing efficiency and therefore requires normalisation. In general, normalisation methods are designed to correct raw counts by compensating for technical factors. However, these methods cannot recover the original, unbiased total mRNA content and instead yield compositional data that convey only relative information between genes[5]. Adding a known amount of spike-in transcripts during sample preparation should, in principle, allow to correct capture and sequencing biases and the recovery of total mRNA content. In practice, incorporating spike-ins into high-throughput protocols (notably droplet-based single-cell isolation) is not widely used, so only a minority of public transcriptomics datasets contain such information.

Here, we leveraged Tabula Muris Senis Smart-seq2 scRNA-seq data, which include ERCC spike-ins[6], and quantified gene expression profiles across a wide variety of cell types throughout the mouse lifespan. Using spike-in normalisation, we find that total mRNA levels vary with age in most cell types analysed. We observe the following trend: most immune cell types show an increase in total mRNA content, whereas most non-immune cell types show a decrease. The number of expressed genes decreased with age in most cell types. We further observed overlap between genes downregulated during ageing and those downregulated during proliferation arrest, suggesting shared regulatory mechanisms. Together, these results suggest changes in the regulation of transcription and cell proliferation with age.

## Results

### TMS data processing and proliferative state evaluation

We analysed Smart-seq2 (FACS) scRNA-seq data from the Tabula Muris Senis atlas[6], which includes ERCC spike-ins. After preprocessing and quality control (SFig. 1), we retained 67390 cells across 21 tissues and 60 tissue-cell types from 3-, 18-, and 24-month-old mice. Of note, 23 of the 60 tissue-cell types were immune cell types (38%). Raw counts were normalised either using ERCC spike-ins or library-size–based total-count scale factors. These approaches were applied in parallel and are hereafter referred to as absolute and global normalisation, as previously described[7].

To assess variation in global transcript abundance, we used the sum of absolute normalised counts per cell as a proxy for total mRNA content. Total mRNA abundance varies with proliferative state[8], and the distribution of cell states captured by single-cell RNA-seq may not reflect their actual in vivo proportions. Consequently, differences in the fraction of proliferating cells across age groups could bias age-related comparisons. To account for this possibility, we classified cells by both cell-cycle stage and proliferative status (quiescent versus proliferating) based on the expression of representative gene sets (Methods).

Most tissues exhibited a low percentage of proliferating cells, with exceptions such as the bone marrow, spleen, and intestine. Additionally, in most cell types, the percentage of proliferating cells remained stable with age, despite fluctuations in some cell types (e.g., BAT_B_cell, Marrow_NK cell, Skin_basal cell, Trachea_T_cell) (SFig. 2a). However, because the proportion of proliferating cells recovered by single-cell RNA-seq may be affected by technical biases in tissue dissociation and cell capture, we did not attempt to further interpret in vivo age-related changes in the abundance of proliferative cells from these data. However, proliferation classification remained useful as an internal validation of our mRNA abundance proxy. Proliferating cells are expected to contain more mRNA and express more genes than quiescent cells. Consistent with this expectation, proliferating cells showed higher total mRNA abundance and more expressed genes than quiescent cells (SFig. 2b–e).

### Age-related changes in total mRNA abundance and number of detected genes

To assess age-associated changes in global gene expression, we fitted a linear model across the three time points for each cell type, for both total mRNA content and the number of genes expressed (including sex and proliferative state as covariates). Among the 60 cell types analysed, 21 exhibited increased total mRNA content, whereas 19 showed a decrease (Fig. 1a). Visual inspection suggested enrichment of immune cell types among those with increased total mRNA content: 13 of the 21 cell types in this group were immune cell types (Fisher’s exact test, p = 0.007, OR = 4.577). By contrast, non-immune cell types were modestly enriched among those with decreased total mRNA content, accounting for 15 of the 19 cell types in this group (p = 0.054, OR = 3.178) (Fig. 1b).

**Figure 1.**
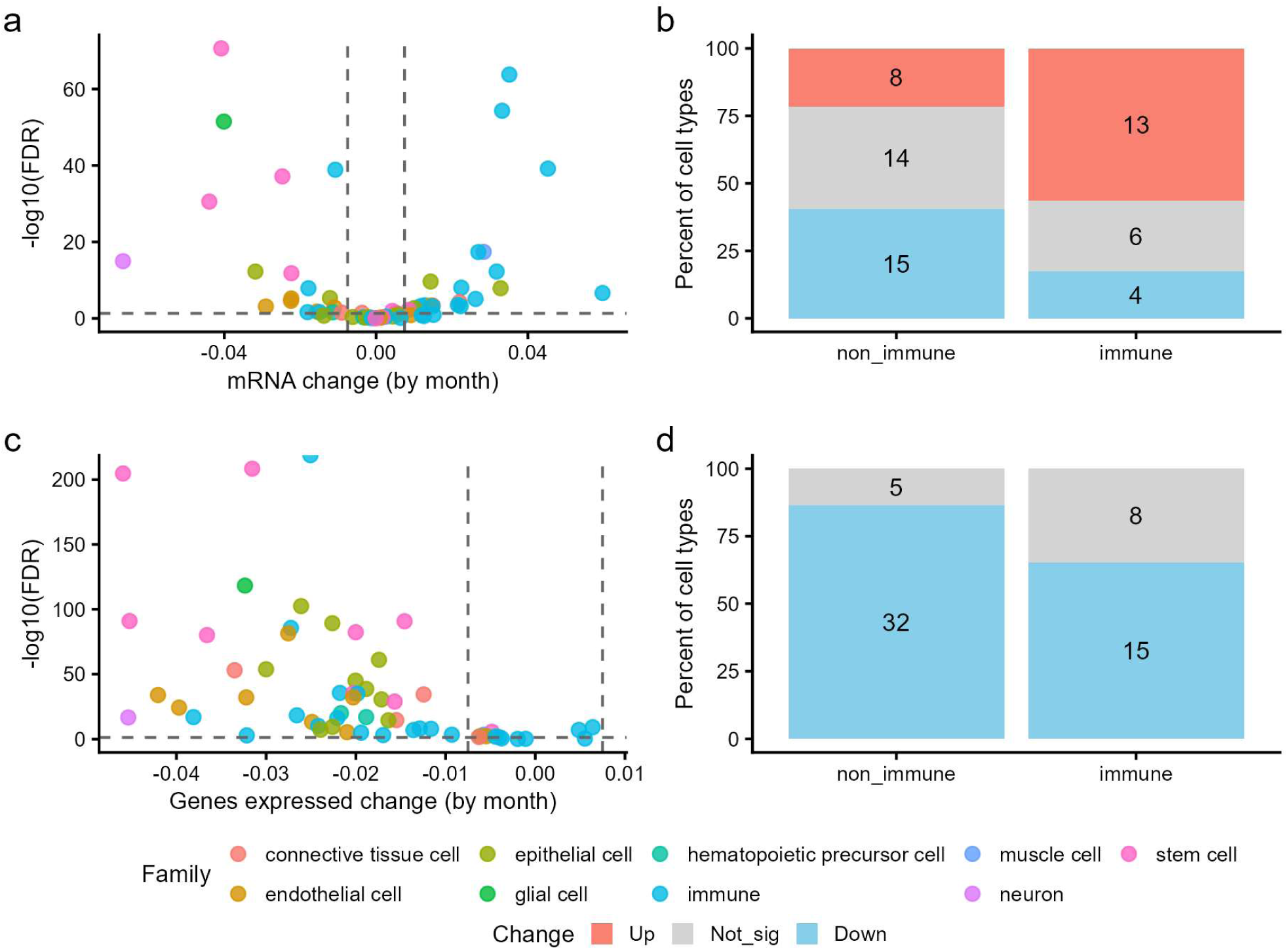
Overall changes in mRNA content during ageing. **a)** Volcano plot of change in mRNA content with age from the linear model (age coefficient). Cells are coloured by the cell type family. **b)** Number of cell types with increase, not changing or decrease in mRNA content from the linear model, separated by immune and non-immune cell types. **c)** Volcano plot of change in genes expressed with age from the linear model (age coefficient). Cells are coloured by the cell type family. **d)** Number of cell types with increase, not changing or decrease in genes expressed from the linear model, separated by immune and non-immune cell types.

More notably, most cell types showed an age-associated reduction in gene count, with decreases observed in 47 cell types (Fig. 1c-d). To help visualise the global gene expression changes with age, we presented the same linear model results as bar plots for mRNA content (SFig. 3a-c) and the number of genes expressed (SFig. 4a-c). In the linear model, in addition to sex, we controlled for proliferative state to minimise the impact of potential technical variability in the proportion of proliferative cells captured. We also tested the linear model without controlling for proliferative state, yielding very similar results for the numbers of cell types with changing mRNA content or the number of genes expressed with age (SFig. 15).

Together, these results point toward a global decline in gene expression activity with age. Even among the cell types showing an increase in total mRNA abundance, the number of detected genes decreased. This pattern is also evident when comparing the linear model coefficients for age-associated changes in total mRNA abundance with those for the number of expressed genes (SFig. 6a). As shown in the next section, this pattern is often driven by a smaller subset of highly expressed genes whose age-associated upregulation dominates total transcript counts. Of note, the reduction in the number of detected genes is unlikely to be due to technical effects, as samples from older ages tended to be sequenced at a deeper level (evaluated by the ERCC counts) (SFig. 6b).

### Differential gene expression analysis with age

Variations in total mRNA abundance arise from the combined effects of changes in individual gene expression. Standard differential expression analyses relying on global normalisation detect shifts in the relative composition of the transcriptome but cannot capture changes in absolute transcript abundance. To characterise gene-level contributions to the age-associated changes described above, we therefore performed differential expression analysis using absolute normalisation.

Linear regression, controlling for sex and proliferating status (Methods), identified cell-type–specific absolute DEGs, with numbers varying across three orders of magnitude across cell types (Fig. 2a-b). Out of the 37 non-immune cell types tested, 32 had more down-regulated than up-regulated absolute DEGs. Of the 23 immune cell types tested, 13 had more down-regulated than up-regulated absolute DEGs. Some cell types showed increased mRNA abundance but had more down-regulated genes. Comparing the mean expression and fold change of the transcriptome in these cell types revealed that the up-regulation of a few highly expressed genes dominates changes in total mRNA abundance (e.g. including Nfkbia, Hspa5, Nr4a1, H2-D1, Nbl1 in Bladder_mesen. Cell and Oaz1, Rps3, Cfl1, Arpc1b, Pfn1, in GAT_myelo. leuko.), despite a larger number of lowly expressed, down-regulated genes (SFig. 7).

**Figure 2.**
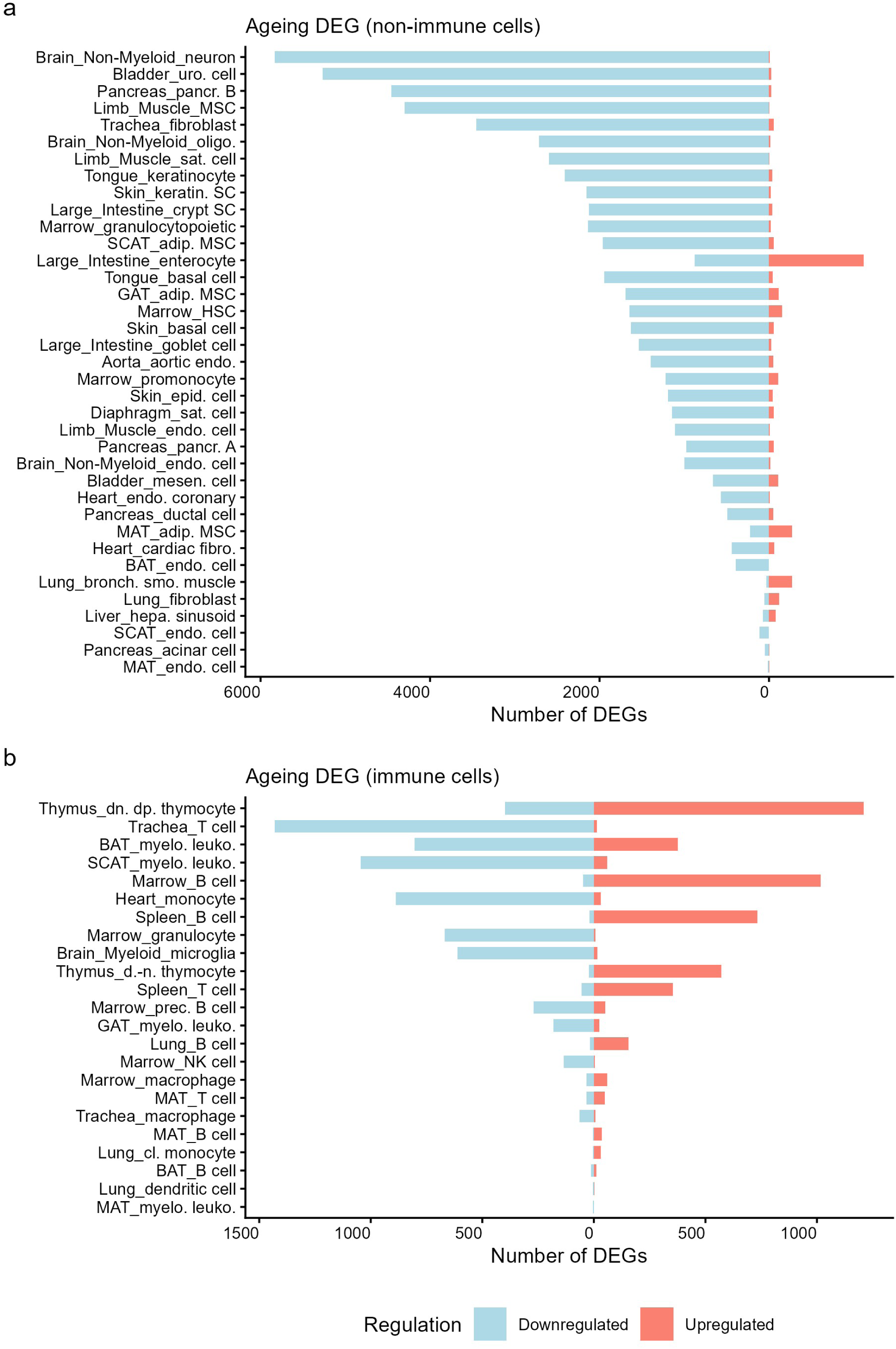
Differential gene expression analysis with age. **a)** Number of DEGs (differentially expressed genes) with age using absolute scaling, from the linear model controlling for sex and proliferative state, by cell type in non-immune cells. Number of DGEs are discriminated by downregulated (left bars in light blue) and upregulated (right bars in light red). Cell types are ordered by the ascending number of total DEGs. **b)** Number of DEGs (differentially expressed genes) with age using absolute scaling, from the linear model controlling for sex and proliferative state, by cell type in immune cells. Number of DGEs are discriminated by downregulated (left bars in light blue) and upregulated (right bars in light red). Cell types are ordered by the ascending number of total DEGs

Since performing differential expression with absolute normalisation is less common than with global normalisation, we thought to compare both methods using the same linear model and the same significance and effect size thresholds. We quantified the number of DEGs that were common to both approaches and those unique to each. Across most cell types, the majority of DEGs were detected by both methods, while absolute normalisation consistently yielded more unique DEGs than global normalisation (SFig. 8a-b). These results are consistent with our previous observation that total mRNA abundance changes with age. Indeed, global normalisation forces samples with potentially different mRNA abundance to have identical total normalised expression. As a thought experiment, if the expression of all the genes were to consistently decrease by exactly 50% across conditions, differential expression with global normalisation would not detect any compositional change (no DEG called), whereas absolute normalisation would lead to all genes being differentially expressed. Although not as extreme, this can be illustrated from our results by comparing the fold-changes obtained by the two methods on three representative examples: a cell type that shows a decrease in total mRNA content (SFig. 9a), a cell type with stable mRNA content (SFig. 9b) and a cell type with increased mRNA content (SFig. 9c). Shifts in log-fold changes are visible as total mRNA abundance varies.

Overall, using absolute scaling, we identified several DEGs across cell types, with a common pattern: more genes are downregulated than upregulated in most non-immune cell types. Some cell types, particularly immune cell types, show the opposite trend, with more genes upregulated. In general, most DEGs were detected by both normalisation methods, as expected; absolute normalisation typically identifies more DEGs. Global normalisation causes a systematic shift in the estimated fold changes in relation to absolute scaling. This comparison emphasises that the choice of normalisation can influence both the magnitude of fold changes and the genes classified as DEGs. Therefore, DEGs obtained using different normalisation approaches should be interpreted with caution.

### Common absolute DEGs associated with ageing

To identify broadly dysregulated processes associated with ageing, we searched for absolute DEGs shared across multiple tissue–cell types. Given the distinct patterns observed between immune and non-immune cell types in the previous analysis, these groups were analysed separately. For non-immune cell types, genes were considered commonly dysregulated if they were identified as absolute DEGs in more than half of the analysed cell types (19 out of 37). Under this criterion, only six genes were consistently upregulated: *Rpl13a, Cfl1, mt-Co2, Tmsb10, Snrpc, and Ptms*. On the other hand, 394 genes were consistently downregulated. Gene Ontology functional enrichment analysis of these downregulated genes (Methods) identified biological processes such as *mRNA processing*, *RNA splicing*, and *regulation of cellular component size*, and molecular functions such as *mRNA binding*, *actin filament binding*, and *microtubule plus-end binding* (Fig. 3a and SFig. 10a). Transcription factor (TF) enrichment analysis using regulons from ENCODE and CHEA (consensus) with enricher[9] revealed potential regulators of these changes, including *TAF1, UBTF, CREB1*, and *YY1* (Fig 3b). Using the CHEA_2022 regulons (Method), we identified additional potential TFs, including *CREM, KDM5b*, and *AF4* (SFig. S10b).

**Figure 3.**
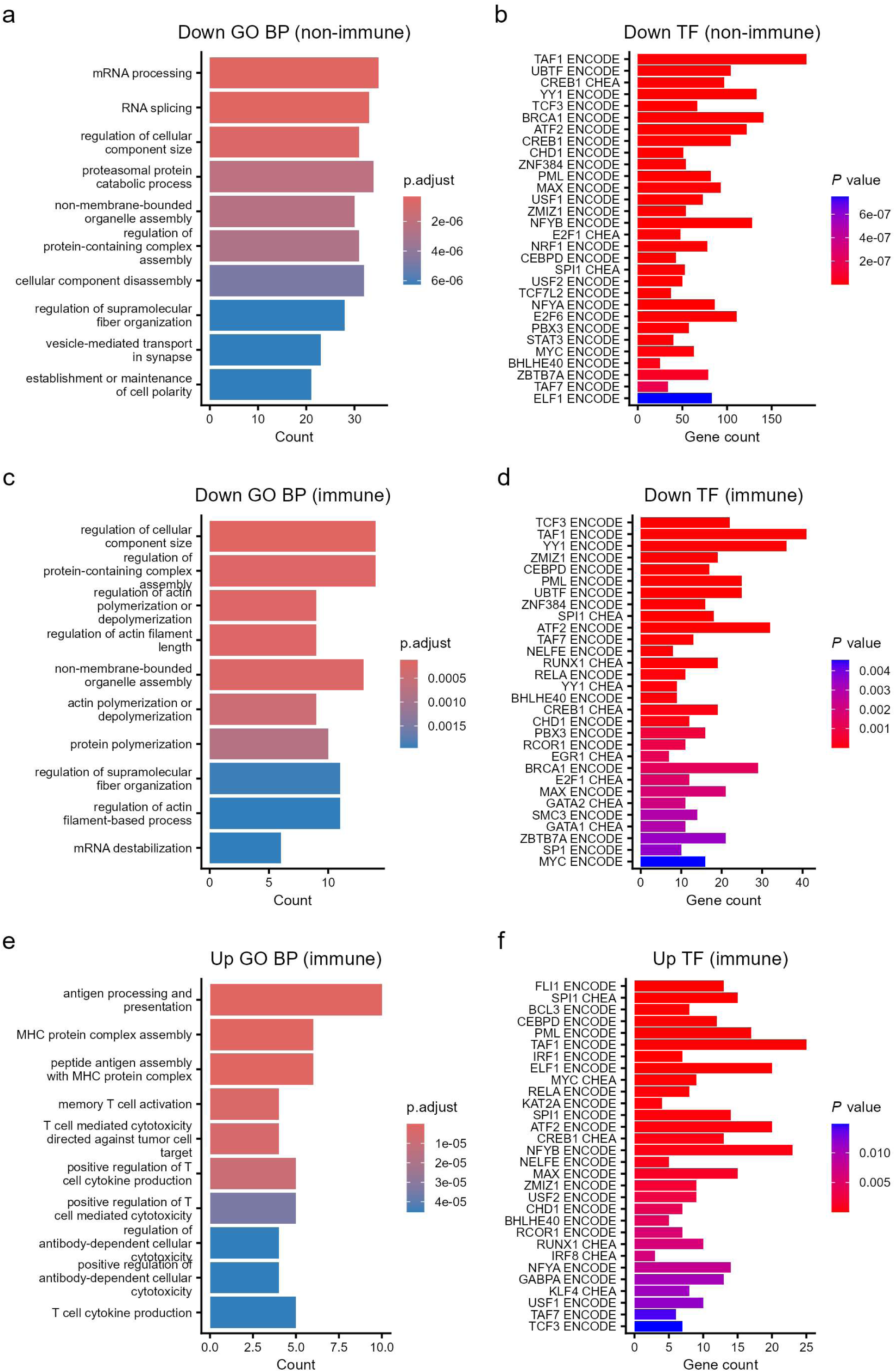
Functional enrichment of gene commonly changing with age. **a)** GO terms functional enrichment at biological process in non-immune cells downregulated genes. **b)** Transcription factors enrichment analysis from enricher with ENCODE and ChEA consensus regulons in non-immune cells downregulated genes. **c)** GO terms functional enrichment at biological process in immune cells downregulated genes. **d)** Transcription factors enrichment analysis from enricher with ENCODE and ChEA consensus regulons in immune cells downregulated genes. **e)** GO terms functional enrichment at biological process in immune cells upregulated genes. **f)** Transcription factors enrichment analysis from enricher with ENCODE and ChEA consensus regulons in immune cells upregulated genes.

For immune cell types, genes were considered commonly dysregulated if they were identified as absolute DEGs in more than one quarter of the cell types (6 out of 23). This one-quarter criterion is based on the observation that roughly half of the cell types are dominated by downregulation, and the other half by upregulation. This criterion revealed 103 common downregulated absolute DEGs functionally enriched for biological processes such as *regulation of cellular component size*, *regulation of protein containing complex* and *regulation of actin polymerization* and *depolymerization* (Fig. 3c), and molecular functions such as *actin filament binding*, and *actin binding* (close to significance level) (SFig. 10c). Enriched transcription factors include *TCF3, TAF1* and *YY1* (SFig. 3d) and, when using ChEA_2022, *AF4, ENL* and *CREM* (SFig. 10d). This shows a similar patterns of downregulation between immune and non-immune cells. Regarding up-regulation in immune cell types, 62 genes were identified as common up-regulated DEGs and were functionally enriched for biological processes such as *antigen processing and presentation*, *MHC protein complex assembly*, and *peptide-antigen assembly with MHC protein complex* (Fig. 3e), and molecular functions including *peptide antigen binding*, *antigen binding*, and *TAP complex binding* (SFig. 10e). Enriched transcription factors include *SPI1, FLI1* and *BCL3* (Fig. 3f) and, when using ChEA_2022, *EKLF*, *TAL1* and *XRN2* (SFig. 10f).

Overall, genes commonly downregulated across multiple cell types may reflect age-related alterations in a shared biological process. The enriched biological functions in common genes downregulated in non-immune cells point to roles in protein and RNA processing, consistent with a general decline in core cellular functions and biosynthetic activity during ageing. Similar functions were observed in the set of genes commonly downregulated in immune cells. The upregulated genes in immune cells were enriched for immune-related functions. In addition, the transcription factors found to have targets enriched among these common DEGs suggest that they may contribute to the coordinated reduction in gene expression. The involvement of these regulatory transcription factors in ageing-related changes could be further investigated in future studies.

### Proliferation-related differential expression analysis

Functional enrichment of the common downregulated absolute DEGs points to reduced cellular biosynthetic activity, including decreased expression of genes involved in RNA processing and protein transport. In addition, several of the enriched transcription factors are known to be involved in proliferation (e.g., *MYC* and *E2F1*). This prompted us to assess the similarity between ageing-related and proliferation-related gene expression changes.

To identify proliferation-related DEGs, we used the same linear model used above to identify age-related DEGs, as it already includes sex and proliferation state as covariates. We note that proliferation states were derived from gene expression data using AUCell with a prior list of proliferating genes. The proliferation states and the differential linear model are therefore not statistically independent. Still, we used the linear model to identify additional genes absent from the proliferation list that may be differentially expressed relative to proliferation. We considered only cell types with at least 50 proliferating and 50 quiescent cells in the proliferation DEG analysis. The number of proliferation-associated differentially expressed genes (DEGs) varied widely, ranging by three orders of magnitude.

To identify genes consistently downregulated upon cessation of proliferation, we defined common downregulated genes as those detected in more than 10 cell types. We selected this threshold to account for the variability in DEG counts, as several cell types exhibited few downregulated genes under our criteria. Notably, we observed more downregulated than upregulated genes in cells transitioning from proliferative to quiescent states (Fig. 4a). This aligns with our observation that quiescent cells exhibit lower mRNA abundance and fewer expressed genes than proliferating cells. Functional enrichment of these downregulated absolute DEGs revealed biological processes such as *ribonucleoprotein complex biogenesis*, *chromosome segregation*, and *RNA splicing* (Fig. 4b), and molecular functions such as *structural component of ribosome*, *mRNA binding*, and *ribonucleoprotein complex binding* (SFig. 11a). Transcription factor enrichment of these same downregulated genes, associated with proliferation arrest, identified regulators such as *TAF1, E2F4, MYC, MAX, YY1*, *BRCA1*, and *ATF2* (Fig. 4c, SFig. 11b). Some of these transcription factors are indeed known to be associated with proliferation (e.g., *E2F1, E2F4, Myc*), and the downregulation of their regulons is consistent with cells transitioning from proliferative to a quiescent state.

**Figure 4.**
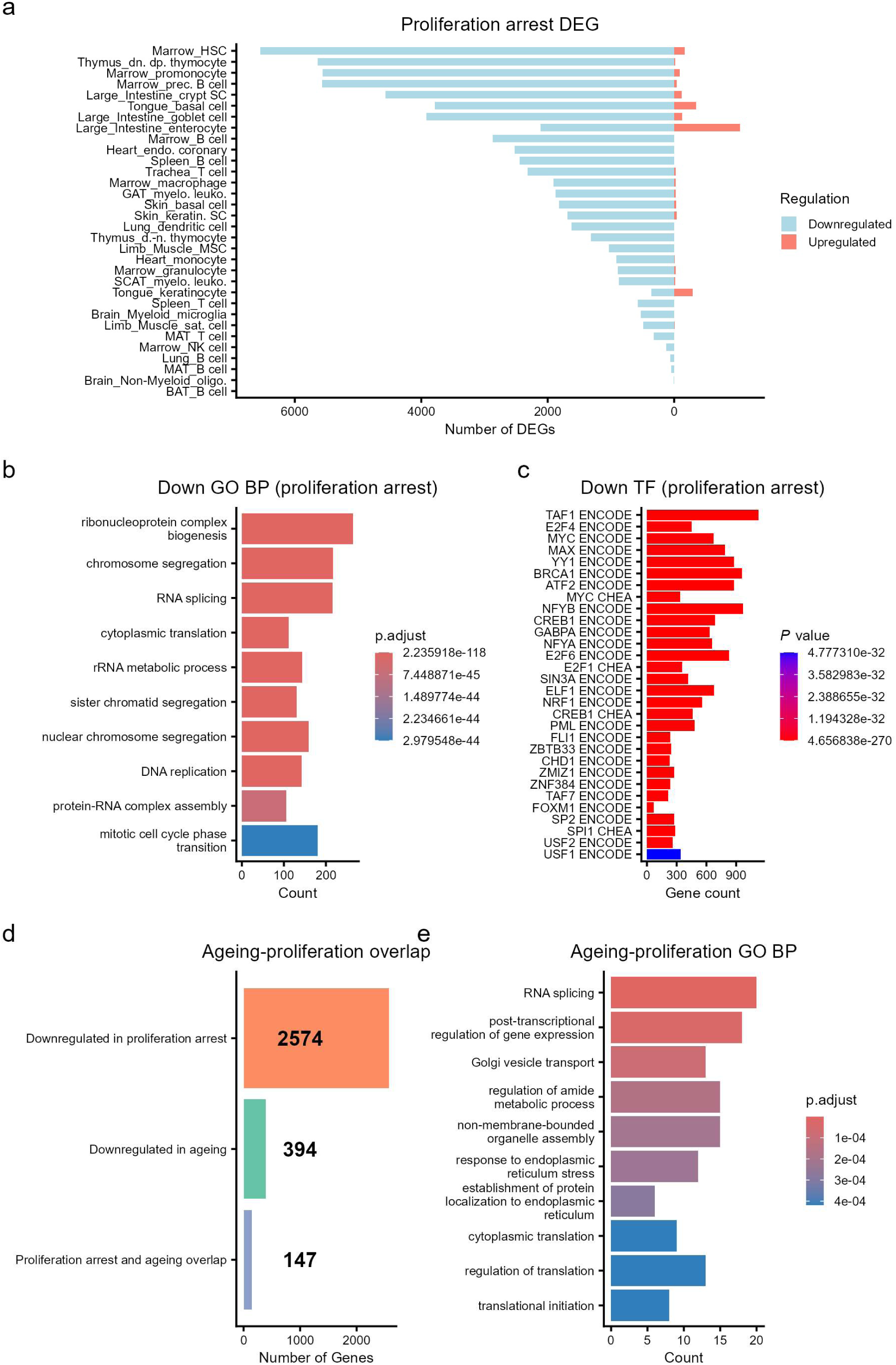
Gene expression associated with cessation of proliferation. **a)** Number of DEGs (differentially expressed genes) with proliferation cessation using absolute scaling, from the linear model controlling for sex and age, by cell type. Number of DGEs are discriminated by downregulated (left bars in light blue) and upregulated (right bars in light red). Cell types are ordered by the ascending number of total DEGs. Note the direction of change is from proliferative to quiescent – proxy of proliferation cessation. **b)** GO terms functional enrichment at biological process of common downregulated genes from proliferation cessation. **c)** Transcription factors enrichment analysis from enricher with ENCODE and ChEA consensus regulons in common downregulated genes from proliferation cessation. **d)** Number of genes common downregulated in proliferation cessation, common downregulated during ageing and then number of overlap genes that are downregulated in several cell types with age in non-immune cells with genes downregulated in the proliferation cessation genes downregulated in both processes. **e)** GO terms functional enrichment at biological process of the overlap genes that are downregulated in several cell types with age in non-immune cells with genes downregulated in the proliferation cessation

We compared genes commonly downregulated with ageing in several non-immune cell types to those downregulated upon cessation of cell proliferation. Over one hundred genes overlapped (Fig. 4d), a higher number than would be observed by chance (random bootstrap, p = 9.999e-05) (SFig. 11c). The genes that overlapped between common ageing downregulated and proliferation arrest downregulated had functional enrichments including *RNA splicing, post-transcriptional regulation of gene expression and Golgi vesicle transpor*t at biological process (Fig. 4e) and *mRNA binding, ribonucleoprotein complex binding and pre-mRNA binding* at molecular function level (SFig. 11d). Additionally, TFs enriched include TAF1, YY1, ATF2, MYC and CREB1 (SFig. 11e-f). Altogether, those overlapping genes appear to be involved in gene expression and cellular biosynthetic activity. Those genes that are downregulated in multiple cell types during ageing and during the transition from proliferation to quiescence could represent a link between growth arrest and ageing.

## Discussion

We leveraged the presence of ERCC spike-ins in the single-cell Tabula Muris Senis Smart-seq2 data to investigate how mRNA abundance varies with age across cell types. Several non-immune cell types showed decreases in total mRNA content and in the number of detected genes per cell, whereas most immune cell types showed increases in total mRNA content. Notably, most cell types exhibiting increased total mRNA showed a concurrent reduction in the number of detected genes, indicating a redistribution of transcript abundance toward fewer, highly expressed genes. Consistent with these patterns, differential expression analysis revealed a predominance of downregulated genes in non-immune cell types, whereas immune cell types showed more balanced responses, with both up- and down-regulation observed. Overall, these results indicate a widespread decline in transcriptome complexity and abundance with age, modulated by immune- and inflammation-associated responses.

Focusing on genes differentially expressed across tissue cell types revealed shared age-related dysregulations. In both non-immune and immune cells, downregulated genes suggested a general age-related reduction in biosynthetic activity, based on functions associated to protein and organelle processing. Conversely, upregulated genes in immune cells were associated with diverse immune functions. Transcription factor enrichment of genes downregulated with age identified regulons, including *TAF1* (core TFIID subunit) [10]; *UBTF* (involved in pol I/II transcription)[11,12]; and the multifunctional *YY1*[13], suggesting a broad age-related decline in transcriptional activity. However, these results are limited by inherent biases toward well-characterised TFs and incomplete regulon data. Furthermore, these correlative associations require experimental validation to establish causality in ageing.

In this study, absolute normalisation was essential to interpret gene-specific effects in the context of global transcriptional decline. An excess of downregulated genes has been reported in other ageing studies using global normalisation, including Tabula Muris Senis[14], multiple human brain cell types[15], and human skeletal muscle[16]. While such asymmetry has been interpreted as evidence for widespread transcriptional repression, under relative normalisation it primarily reflects compositional changes and does not, by itself, imply a decrease in total mRNA content. Taken together, however, results from both absolute and global normalisation suggest that ageing is characterised by a predominance of gene downregulation at both the absolute and relative levels, at least in non-immune cell types. Additionally, evidence indicates that mRNA translation rates decline with age across multiple species, suggesting reduced protein production[17], and a meta-analysis of ageing transcriptomes has proposed a decrease in transcription [3]. Together, these findings are consistent with a general decline in cellular activity with age in several non-immune cell types.

At a functional level, it is tempting to speculate that the observed global reduction in gene expression is associated with age-related tissue atrophy. For example, the brain is known to undergo atrophy with age[18] and, correspondingly, we observed reduced global gene expression in neurons, oligodendrocytes, microglia, and brain endothelial cells. Similarly, we detected decreased global gene expression in mesenchymal stem cells, satellite cells, and endothelial cells in limb muscles, which may contribute to the well-documented age-related loss of muscle mass (sarcopenia) [19].

It will be important to elucidate the molecular mechanisms underlying these changes. For example, increased RNA polymerase II elongation speed with age has been reported[20]. This does not necessarily conflict with the widespread reduction in mRNA content observed here, as elongation is only one of several parameters governing gene expression dynamics. Changes in transcriptional burst frequency and size, as well as in mRNA degradation rates [21] and translation rates, are also likely to contribute [17]. Resolving how these processes jointly shape gene expression and understanding distinct observations altogether will require systematic mechanistic studies across cell types and ages.

We noted a parallel between ageing and cessation of proliferation, both marked by a predominance of gene downregulation, shared enriched TFs (e.g., *Myc* and *E2F1*), and similar functional enrichment, suggesting decreased cell biosynthetic activity. To further explore the link between ageing and the cessation of proliferation, we compared genes commonly downregulated in non-immune cells with those downregulated during growth arrest. We identified 147 genes shared between these two groups. These similarities suggest that signalling pathways that regulate cell proliferation may also contribute to the global downregulation of ageing-associated genes. Future studies should explore this hypothesis experimentally.

Ageing theories generally fall into two categories: developmental and stochastic damage models[22]. The widespread gene downregulation with age could align with both. For instance, DNA damage could affect transcription[23], and was associated with RNA polymerase stalling during ageing[24]. Alternatively, the decline in mRNA content with age may reflect a continuation of developmental programs into adulthood, marked by reduced growth and renewal. While both models are consistent with our observations, we suggest that reduced mRNA content may reflect a developmental trajectory. Accordingly, activated embryonic and adult stem/progenitor cells have been reported to undergo hypertranscription (a global increase in mRNA content) during development, organ maintenance, and regeneration[7]. High transcriptional diversity, corresponding to a large number of distinct genes expressed, is also characteristic of stem cells and decreases during differentiation [25]. In contrast, our observation that many cell types exhibit reduced mRNA content and fewer detected genes with age points to widespread reductions in gene expression (hypotranscription). One hypothesis is that decreased mRNA content may contribute to impaired organ maintenance/regeneration and to the onset and/or progression of ageing phenotypes. Hypertranscription has indeed been recognised as a feature of aggressive tumours and is associated with a worse prognosis across cancers[26]. The age-related hypotranscription we observed here further supports the complex relationship, including opposite patterns, between ageing and cancer, previously noted by some of us in human bulk transcriptomics analyses[27]. Additionally, our observation of a global reduction in gene expression with age, together with evidence that suppression of cell proliferation may contribute to this decline, is consistent with a proposed model in which biological processes that evolved to prevent cancer, such as tumour-suppressor mechanisms, inadvertently promote ageing [28,29].

In this work, using absolute scaling, we identified distinct ageing patterns across cell types. Non-immune cells showed a dominant trend of downregulation and reduced mRNA content, with functional declines in RNA and protein processing suggesting diminished biosynthesis. In contrast, immune cells exhibited a more balanced transcriptional profile, with upregulated genes associated with enhanced immune function. Our findings also suggest a link between global downregulation and control of cell proliferation. We hope to encourage the wider adoption of absolute normalisation techniques in future ageing research.

### Limitations

While global normalisation cannot account for total mRNA content, absolute normalisation, which uses external spike-in controls, preserves information about the sample’s initial total mRNA. Despite evidence of adequate performance in single-cell RNA-seq[30], this approach has its own drawbacks, including variation in the amount of spike-in RNA added across samples (pipetting inconsistencies or degradation of spike RNA), which leads to sample-to-sample variation in spike-in levels [31]. Synthetic spike-in transcripts may behave differently from endogenous transcripts in terms of capture efficiency[32], for example was estimated that endogenous transcripts are indeed more efficiently captured than spike-ins by about one order of magnitude[33]. Although not perfect, this approach remains one of the few ways to estimate global transcriptomic variation, which can have important biological consequences[34], notably for ageing [3]. However, it would be relevant to expand this analysis and replicate the results with larger datasets and using distinct spikes or complementary methods.

Although this is a large ageing scRNA-seq dataset with spike-ins, it remains limited in the range of ages, the number of biological replicates, and the coverage of cell types, with some populations represented by relatively few cells. These constraints are particularly relevant for scRNA-seq due to the stochastic nature of gene expression at the cellular level [35], and the stochastic, low-efficiency capture of transcripts [36], among other factors. These features increase the risk of false discoveries. This issue can be mitigated using pseudobulk approaches[37], however, the number of biological replicates was insufficient to support robust pseudobulk aggregation, and we therefore relied on cell-level models. To limit false positives, we applied a linear model with sex and proliferative state as covariates and imposed more stringent effect-size thresholds. Additionally, age-associated changes in mRNA content are most pronounced at 24 months, the oldest age in the dataset. However, this time point includes only male samples. Sex differences in ageing-related gene expression have been reported[38], and studies of frailty suggest that female mice may maintain a longer health span than males[39]. Altogether, those limitations reinforce the need for further work using larger, age-resolved, sex-balanced, and ERCC spike-in datasets to validate and extend these findings, ideally extending this analysis to human datasets.

## Methods

### Retrieving single-cell RNA sequencing data

We downloaded the Tabula Muris Senis (TMS) FACS .h5ad dataset (adata_with_ercc_genecode_counts_for_gatk_with_metadata.h5ad), which includes ERCC spike-in counts, from the czb-tabula-muris-senis S3 repository.

### Handling SingleCellExperiment objects and plotting

The Tabula Muris Senis (TMS) .h5ad file was imported into R and converted to a SingleCellExperiment (SCE) object using the zellkonverter package. The SCE object was processed using SingleCellExperiment, tidySingleCellExperiment, scran, and scater, which are available through Bioconductor[40]. Data analysis was performed in R using base functions and the tidyverse ecosystem. Visualisations were generated with ggplot2.

### QC, filtering and normalisation

Gene biotypes were annotated using biomaRt[41] (mmusculus_gene_ensembl). Only protein-coding features were retained; Malat1 counts were stored separately for quality control. Features detected in fewer than three cells were excluded. Cells lacking a TMS cell identity annotation (cell_ontology_class) were removed, along with cells with fewer than 5,000 counts or fewer than 200 detected genes, consistent with the original TMS thresholds[6].

Cells with less than 0.25% or more than 25% ERCC spike-in content were excluded. Low ERCC percentages may indicate doublets or multiplets[42], particularly homotypic events that are difficult to detect and can bias absolute mRNA estimates upward. High ERCC percentages may indicate cell damage; therefore, an upper threshold was also applied. To further ensure reliable spike-in quantification, cells with fewer than 1,000 ERCC counts or fewer than 10 detected ERCC features were removed. Cells with more than 20% mitochondrial transcripts were excluded. As an additional quality-control step, cells with fewer than 100 Malat1 counts were discarded [43].

To ensure sufficient representation across age groups, we retained only cell types with at least 50 cells at each time point (3, 18, and 24 months). Distributions of counts after filtering are shown in Supplementary Fig. 1, including total mRNA raw and ERCC counts (SFig. 1a), number of detected mRNA and ERCC genes (SFig. 1b), and the relationship between mRNA content (normalized) and gene detection (SFig. 1c). The comparison of the distribution of raw mRNA counts with normalized (absolute scaling) mRNA counts, used as proxy of mRNA content (SFig. 1d) and the distribution of the percentage of ERCC (SFig 1e).

Cell types were defined as combinations of tissue and cell ontology annotations (cell_ontology_class), following the nomenclature previously described[44], with “leukocyte” relabelled as “immune” for simplicity.

Absolute normalisation was performed using ERCC spike-ins[7]. Size factors were estimated per cell using ERCC counts (computeSpikeFactors, scran) and applied to the expression matrix using normalizeCounts (scater), both without and with log transformation (adding a pseudocount of 1). ERCC counts showed a linear relationship with their known input concentrations. For comparison, global normalisation based on total counts was performed using logNormCounts (scater). Potential doublets were identified using scDblFinder[45], with tissues treated as samples and were removed prior to downstream analysis.

### Cell cycle stage and proliferation assignments

Each cell was assigned a cell-cycle stage (G1, S, G2/M) using cyclone[46](scran). In addition, cells were classified as proliferative or quiescent based on a proliferation-associated gene set[47][48]and classified cells with a score bigger than 0.15 as proliferative; all others were classified as quiescent. This definition does not distinguish between reversible quiescence and terminal differentiation.

Most cells were classified as quiescent and were in G1 phase, consistent with the adult and aged nature of the samples. The distribution of cell-cycle stages across proliferative and quiescent groups was concordant (SFig. 1f): quiescent cells were predominantly in G1, whereas proliferative cells were distributed across G1, S, and G2/M. Only a small number of discrepancies were observed between the two classification approaches (e.g., quiescent cells assigned to S or G2/M).

### mRNA content and number of expressed genes

Total mRNA content was approximated as the sum of absolute normalised gene counts per cell. For each cell type, mRNA content and the number of expressed genes (in log2 scale) were analysed using linear models (R lm function) with age treated as a continuous variable, and sex and proliferative state included as covariates. Cell types were classified as age-dependent using thresholds of an effect size > 0.0075 (log fold change per month, ∼10% from 3m to 24m) and a false discovery rate (FDR) <0.05. Enrichment of immune cell types among those with increased mRNA content, and of non-immune cell types among those with decreased mRNA content, was assessed using Fisher’s exact test in R.

### Differential expression analysis

Differential expression with age was assessed separately for each cell type. To model age as a continuous variable, we used a linear model implemented in testLinearModel (scran), including sex and proliferative state as covariates. Genes were considered differentially expressed if the age coefficient was ≥0.05 (log fold change per month) and the false discovery rate (FDR) was <0.01. Analyses were performed using both absolute and global normalisation for comparison; unless stated otherwise, downstream analyses are based on absolute normalisation.

To compare proliferative and quiescent cells, we used the same linear framework but focused on the coefficient associated with proliferative status. Genes were considered differentially expressed if |logFC| ≥ 0.5 and FDR < 0.01. Only cell types with at least 50 cells in each group (proliferative and quiescent) were included in this analysis.

### Functional and transcription factors enrichment

Functional enrichment analysis was performed using clusterProfiler[49] for Gene Ontology (GO) biological process and molecular function categories. Enrichment was conducted using genes present in the dataset as the background (universe) and annotations from org.Mm.eg.db. Redundant GO terms were reduced using the simplify function in clusterProfiler.

Transcription factor enrichment was performed using enrichR[50], with the ENCODE_and_ChEA_Consensus_TFs_from_ChIP-X and ChEA_2022 databases. These resources are based on regulons derived from chromatin immunoprecipitation experiments. The ENCODE_and_ChEA_Consensus dataset represents consensus regulons aggregated across multiple studies, whereas ChEA_2022 compiles regulons from individual experiments. Although the use of both introduces some redundancy, it provides broader coverage of potential transcriptional regulators.

### Common genes

To identify genes consistently differentially expressed across cell types, we counted the number of cell types in which each gene was detected as differentially expressed. For non-immune cell types, genes were defined as common DEGs (separately for up- and downregulation) if they were identified in at least half of the cell types.

For immune cell types, the threshold was reduced to one-quarter of the cell type threshold, reflecting a more balanced distribution of up- and downregulated genes across these populations.

For proliferation arrest, we observed that 19 cell types had more than 1,000 downregulated genes, whereas the remaining cell types had relatively few. To avoid bias from sparsely represented signals, we defined common DEGs using a threshold of 10 cell types (approximately half of the strongly affected cell types).

### Code

Full script with package versions, code chunks and parameters will be available through the GitHub repository.

## Acknowledgements

We grateful to the members of the Genomics of Ageing and Rejuvenation Lab, additionally to Ana Pena and Lara Escobar for valuable discussions and suggestions.

During the preparation of this work, the authors used AI-assisted tools, Grammarly, Gemini and ChatGPT, to improve readability and language and improve code. After using these tools, we reviewed and edited the text as needed and take full responsibility for the content of the publication. Work in our lab is supported by grants from LongeCity and the Biotechnology and Biological Sciences Research Council. Idálio de Jesus Viegas was supported by an EMBO fellowship (EMBO ALTF 1028-2022).

## Supplementary

**Figure S1.**
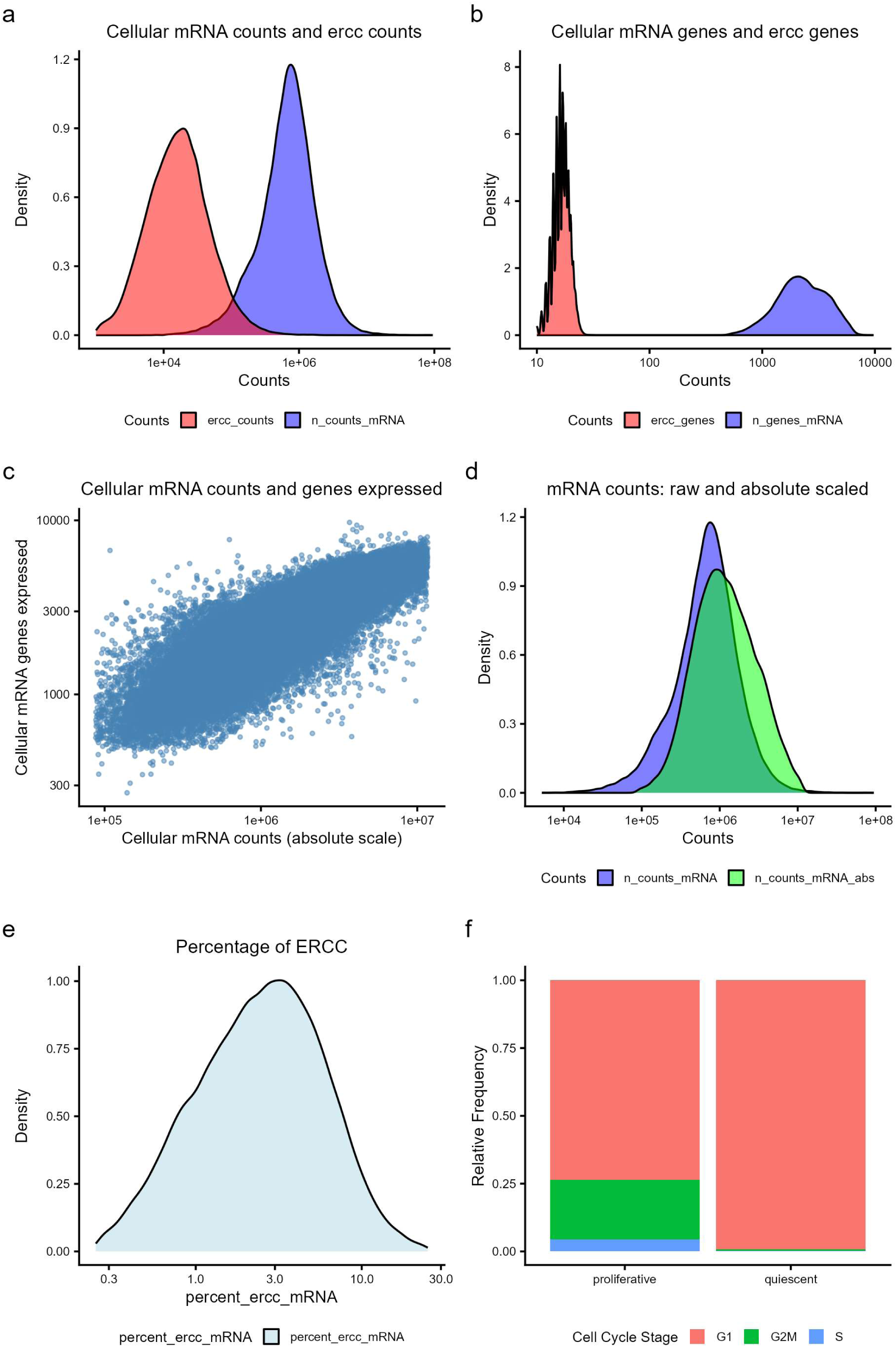
Overview of the dataset after QC filtering. **a)** Cellular counts and ERCC counts. **b)** Cellular genes detected and ERCC detected genes. **c)** mRNA content and number of genes detected with expression **d)** Cellular counts (raw) compared with absolute scaled normalized counts. **e)** percentage of ERCC relative to mRNA. **h)** Comparison of cell cycle stage assignment and proliferation status assignment.

**Figure S2.**
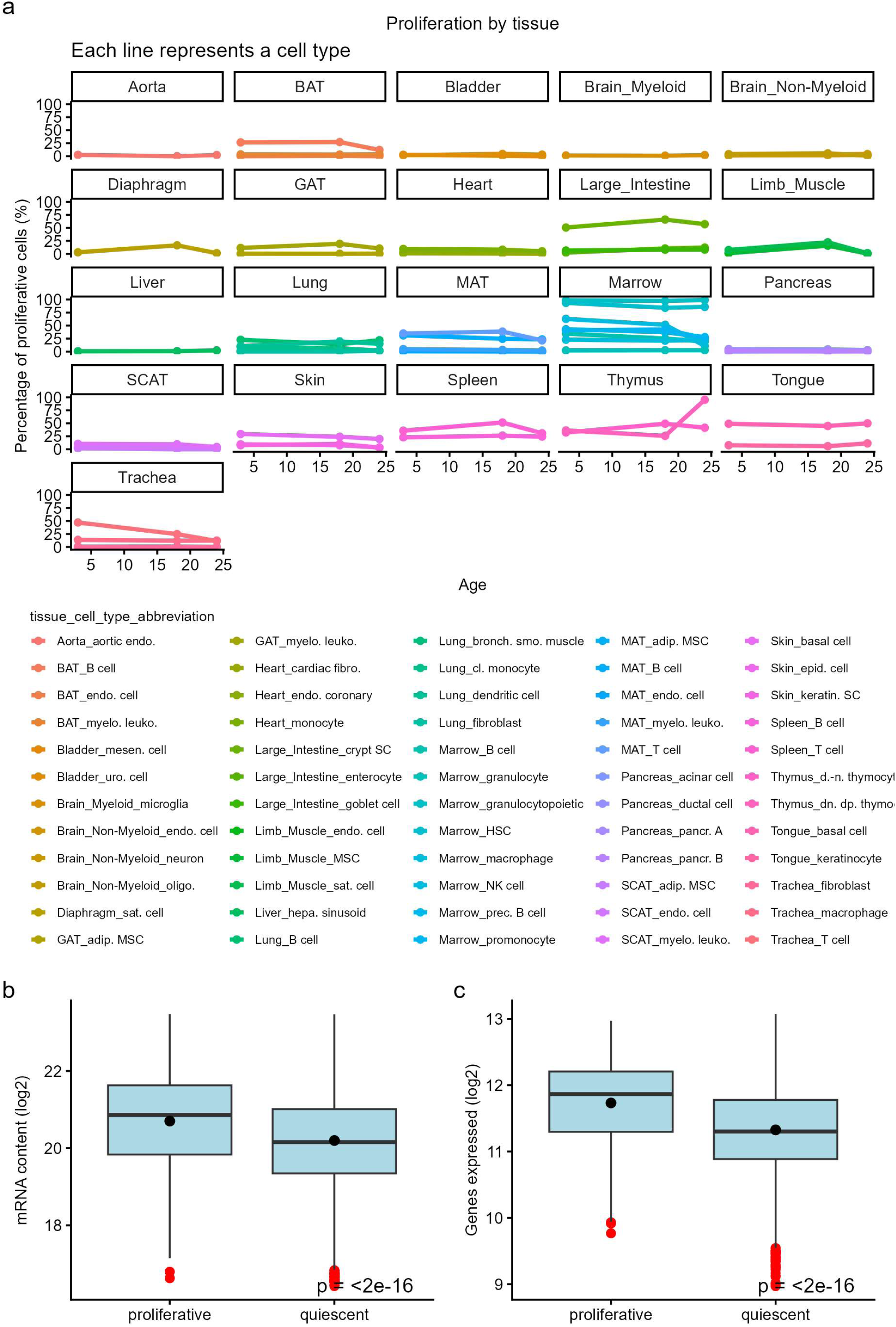
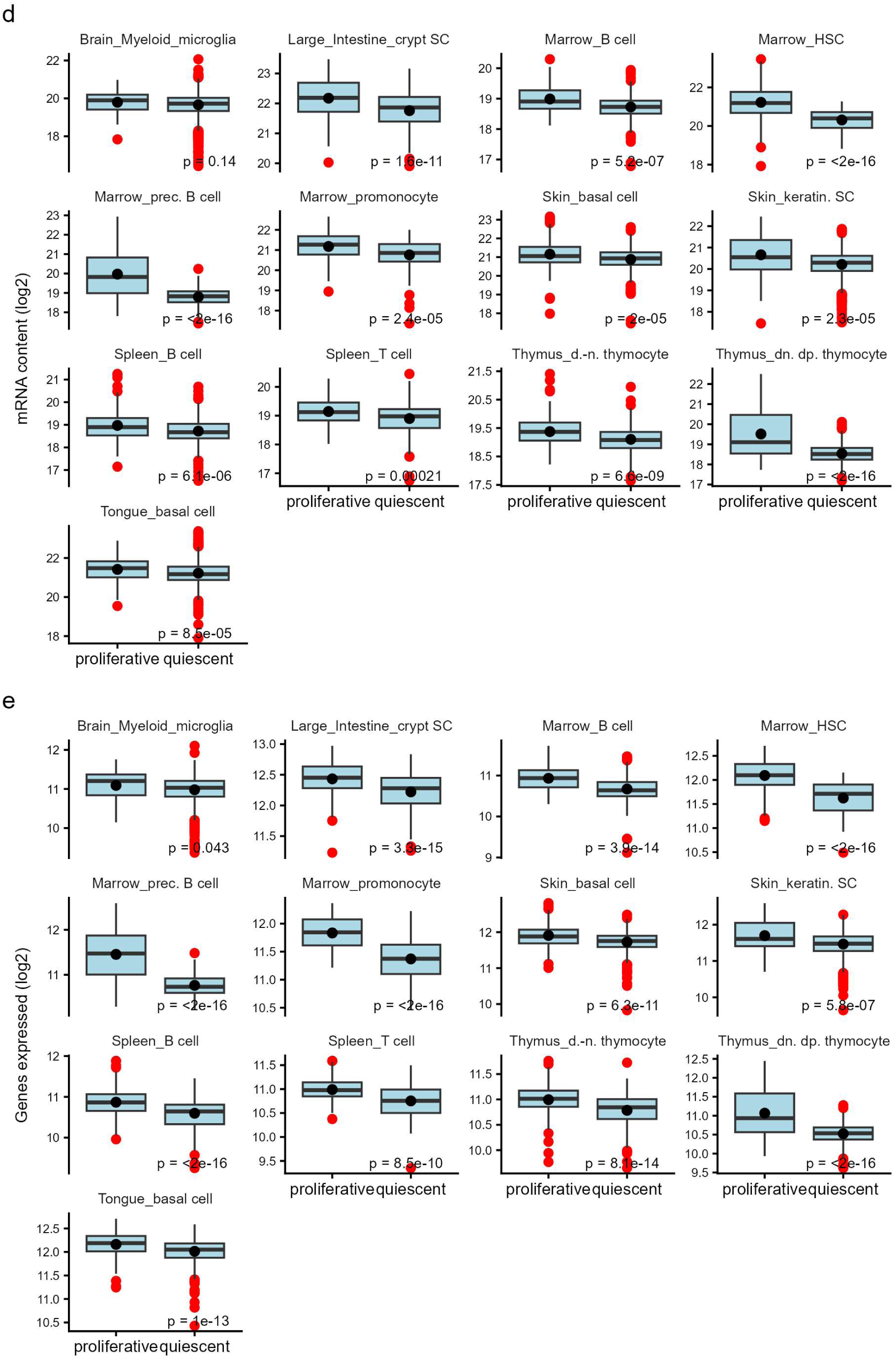
Overall changes in mRNA content and genes expressed with proliferative state. **a)** Percentage of proliferating cells by cell type, in the different tissues, by age in the dataset. **b)** mRNA content (sum of absolute scale normalized counts) in proliferative and quiescent cells, from adult animals (3 months old). P value from a t-test in the bottom right corner. **c)** Number of genes expressed in proliferative and quiescent cells, from adult animals (3 months old). P value from a t-test in the bottom right corner. **d)** mRNA content in proliferative and quiescent cells, from adult animals (3 months old) discriminated by cell type (cell types with at least 50 cells in each group). P value from a t-test in the bottom right corner. **e)** Number of genes expressed in proliferative and quiescent cells, from adult animals (3 months old) discriminated by cell type (cell types with at least 50 cells in each group). P value from a t-test in the bottom right corner.

**Figure S3.**
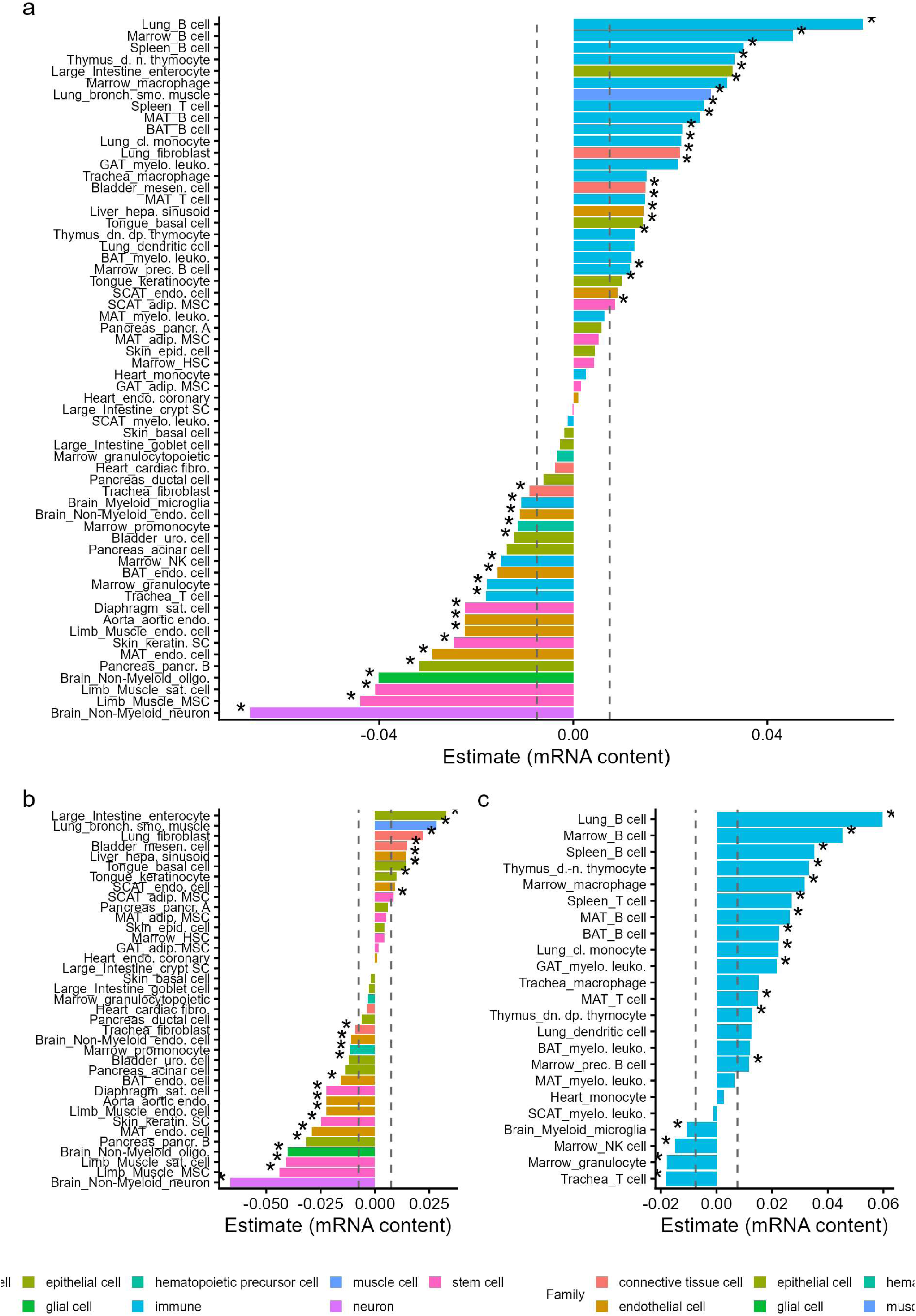
mRNA content change with age from the linear model visualized as a bar plot. **a)** Bar plot of change in mRNA content with age from the linear model (age coefficient). Cells are coloured by the cell type family. * Denotes the cell types that FDR was lower than 0.05 and the vertical lines indicates the selected coefficient threshold. Negative values (left) represent a decrease with age, while positive values (right) represent an increase with age. **b)** Bar plot of change in mRNA content with age from the linear model (age coefficient), the same data as in S3a, but only with non-immune cells. Cells are coloured by the cell type family. * Denotes the cell types that FDR was lower than 0.05 and the vertical lines indicates the selected coefficient threshold. Negative values (left) represent a decrease with age, while positive values (right) represent an increase with age. **c)** Bar plot of change in mRNA content with age from the linear model (age coefficient), the same data as in S3a, but only with immune cells. Cells are coloured by the cell type family. * Denotes the cell types that FDR was lower than 0.05 and the vertical lines indicates the selected coefficient threshold. Negative values (left) represent a decrease with age, while positive values (right) represent an increase with age.

**Figure S4.**
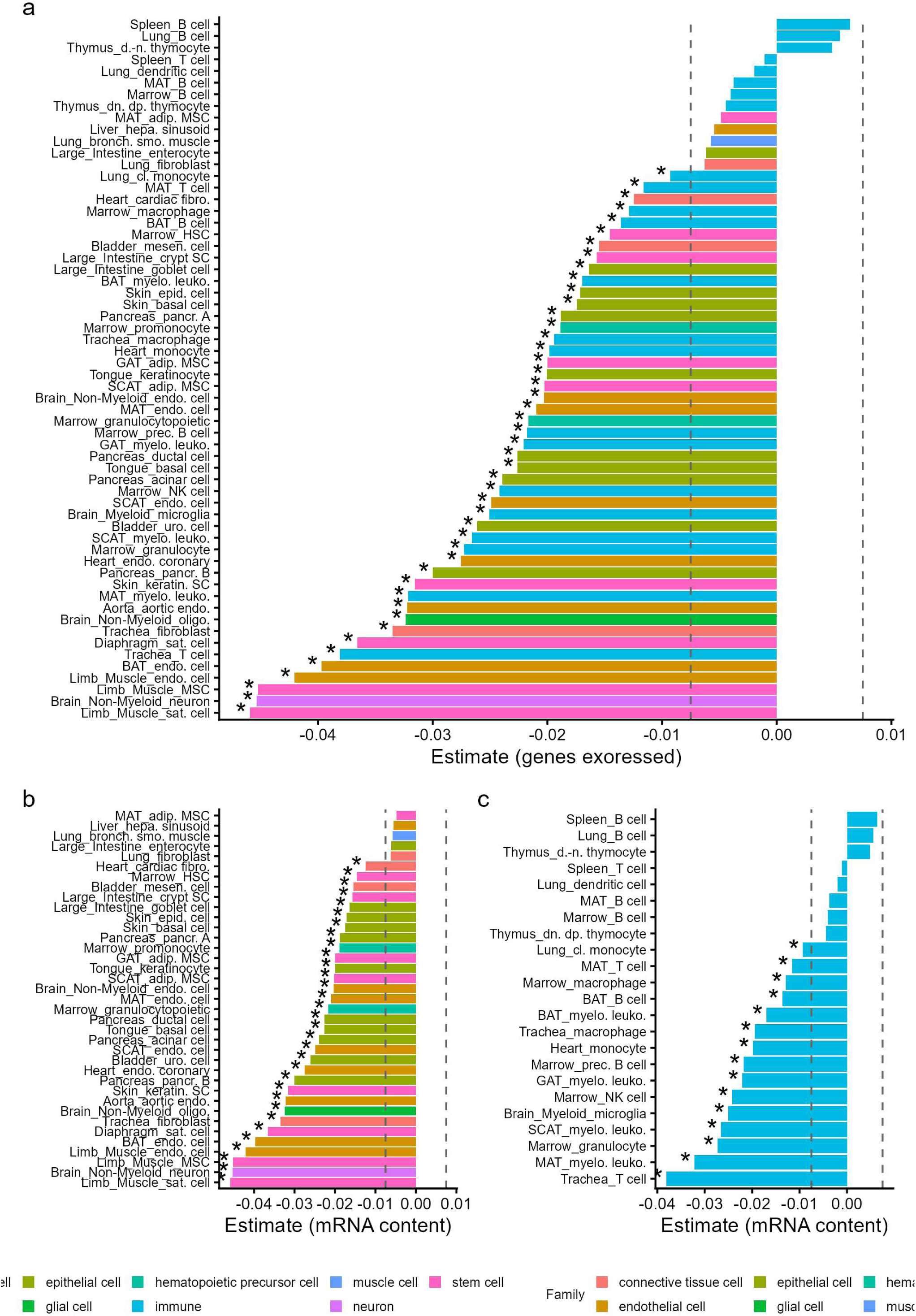
Number of genes expressed change with age from the linear model visualized as a bar plot. **a)** Bar plot of change in genes expressed with age from the linear model (age coefficient). Cells are coloured by the cell type family. * Denotes the cell types that FDR was lower than 0.05 and the vertical lines indicates the selected coefficient threshold. Negative values (left) represent a decrease with age, while positive values (right) represent an increase with age. **b)** Bar plot of change in genes expressed with age from the linear model (age coefficient), the same data as in S3a, but only with non-immune cells. Cells are coloured by the cell type family. * Denotes the cell types that FDR was lower than 0.05 and the vertical lines indicates the selected coefficient threshold. Negative values (left) represent a decrease with age, while positive values (right) represent an increase with age. **c)** Bar plot of change in genes expressed with age from the linear model (age coefficient), the same data as in S3a, but only with immune cells. Cells are coloured by the cell type family. * Denotes the cell types that FDR was lower than 0.05 and the vertical lines indicates the selected coefficient threshold. Negative values (left) represent a decrease with age, while positive values (right) represent an increase with age.

**Figure S5.**
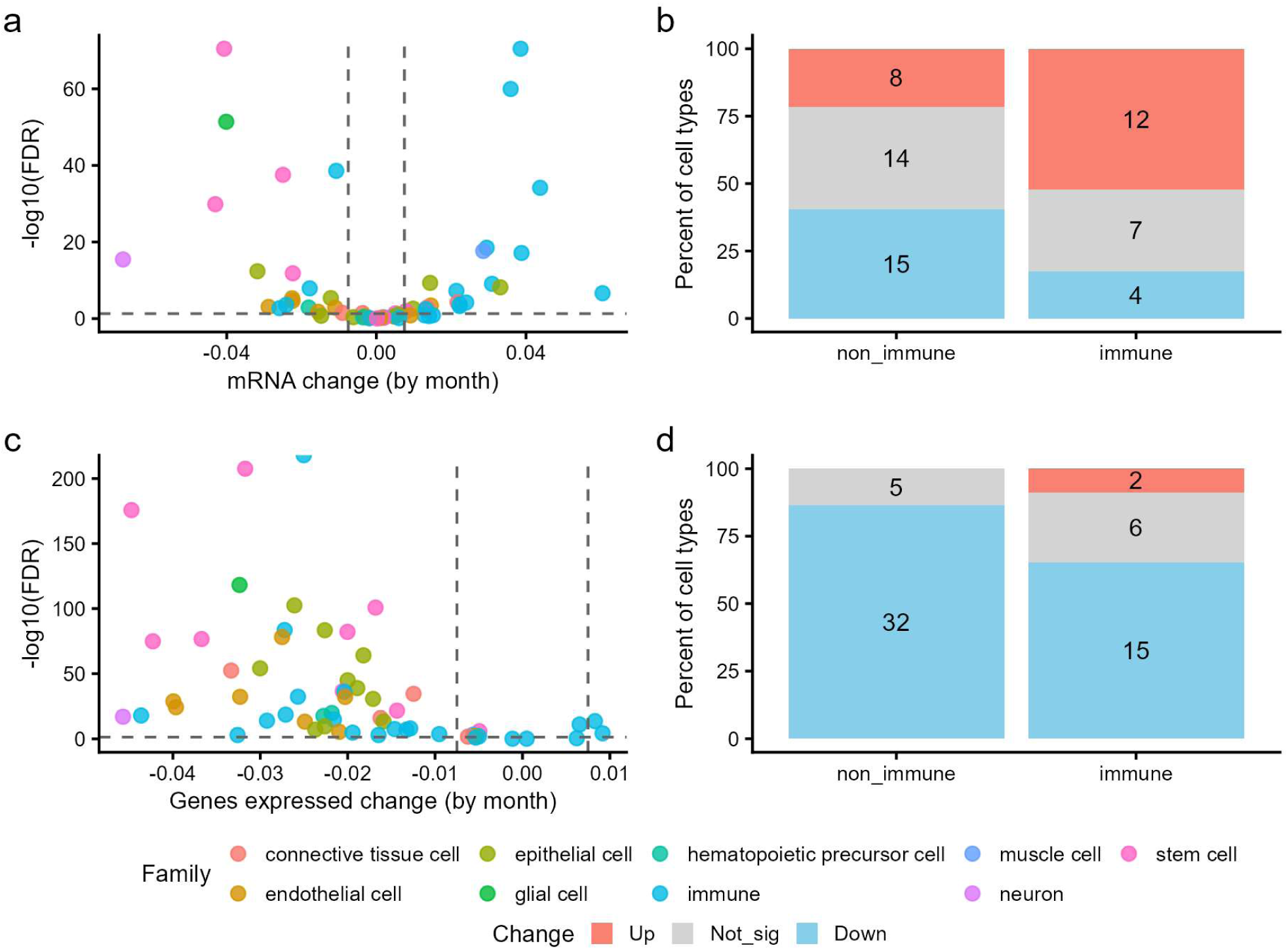
Overall changes in mRNA content during ageing without controlling for proliferative state. **a)** Volcano plot of change in mRNA content with age from the linear model (age coefficient). Cells are coloured by the cell type family. **b)** Number of cell types with increase, not changing or decrease in mRNA content from the linear model, separated by immune and non-immune cell types. **c)** Volcano plot of change in genes expressed with age from the linear model (age coefficient). Cells are coloured by the cell type family. **d)** Number of cell types with increase, not changing or decrease in genes expressed from the linear model, separated by immune and non-immune cell types.

**Figure S6.**
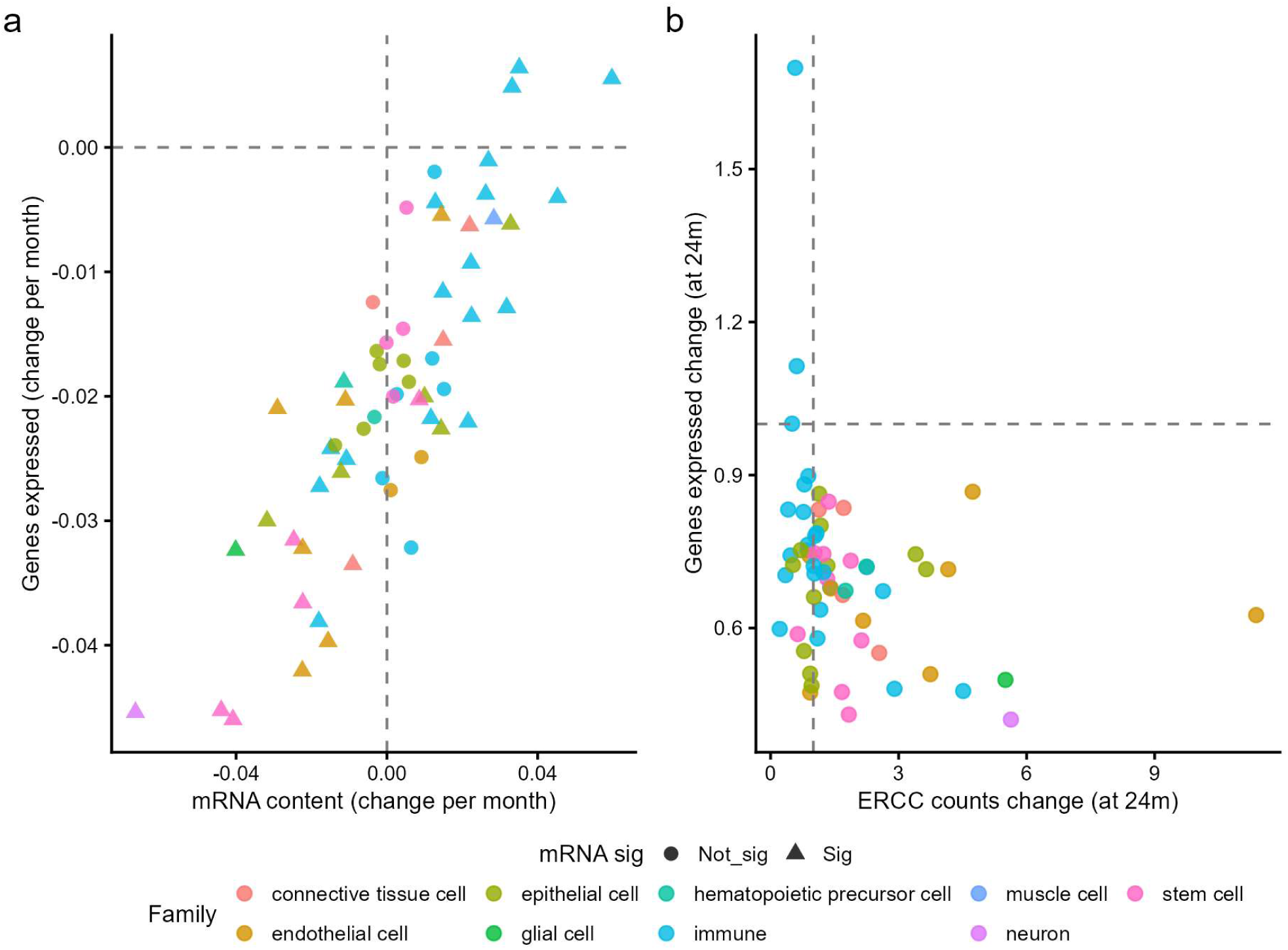
Comparison of linear model estimates for mRNA content and genes expressed. **a)** Comparison of the age associated change in mRNA content (x axe) with the age associated change in genes expressed (y axe). Cells are coloured by family, and the symbols represents significance in the linear model (FDR less than 0.05. **b)** Relation between the number of genes detected with expression (median fold change at 24m) and the sequencing effort, evaluated by ERCC counts (median fold change at 24m). Cell types are labelled by family.

**Figure S7.**
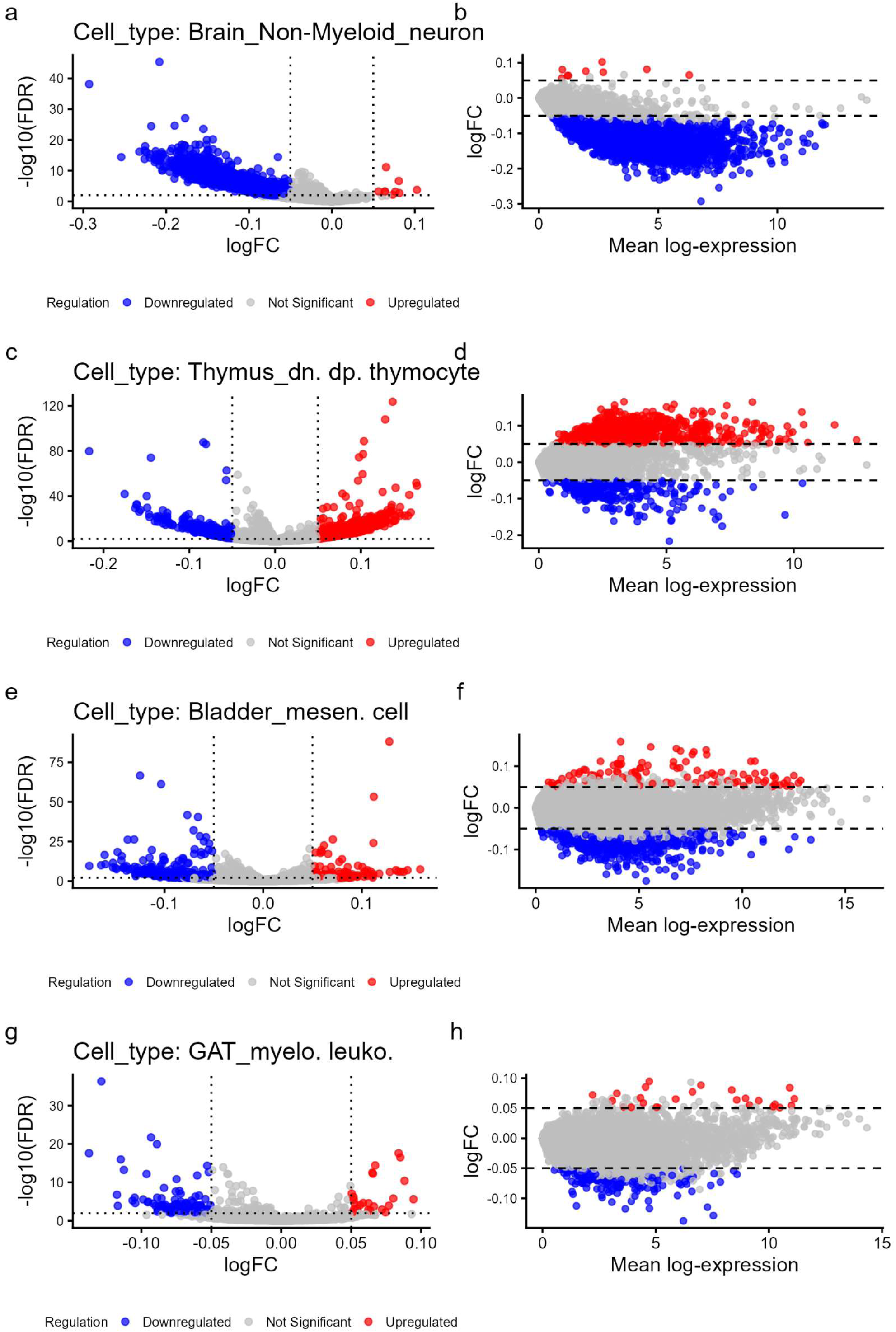
Differential gene expression with age, examples. **a)** Volcano (logFC vs FDR) plot of Brain_Non_mieloid_neuron**. b)** MA plot (mean expression vs logFC) of Brain_Non_mieloid_neuron. **c)** Volcano (logFC vs FDR) plot of Thymus_dn_dp_thymocyte**. d)** MA plot (mean expression vs logFC) of Thymus_dn_dp_thymocyte. e) Volcano (logFC vs FDR) plot of Bladder_mesen.cell**. f)** MA plot (mean expression vs logFC) of Bladder_mesen.cell. **g)** Volcano (logFC vs FDR) plot of GAT_myelo.leuko**. h)** MA plot (mean expression vs logFC) of GAT_myelo.leuko.

**Figure S8.**
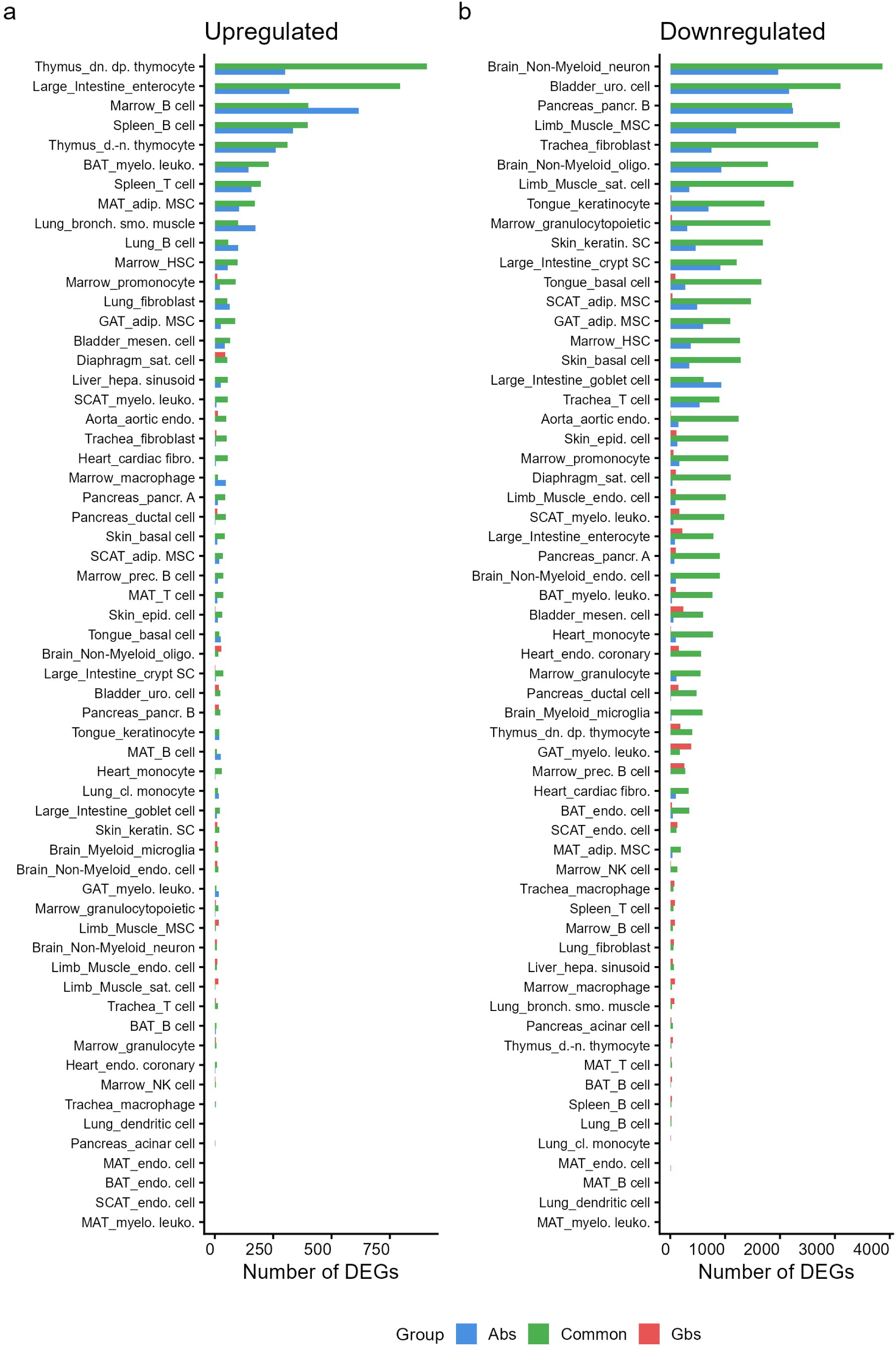
Differential gene expression with age comparison of absolute scaling with global scaling. **a)** Comparison of the upregulated DEGs identified with absolute scaling with DEGs identified with global scaling. Bar coloured by the group (blue the number of genes identified only with absolute scaling, green genes identified both with absolute and global scaling and red genes identified only with global scaling). **b)** Comparison of the downregulated DEGs identified with absolute scaling with DEGs identified with global scaling. Bar coloured by the group (blue the number of genes identified only with absolute scaling, green genes identified both with absolute and global scaling and red genes identified only with global scaling).

**Figure S9.**
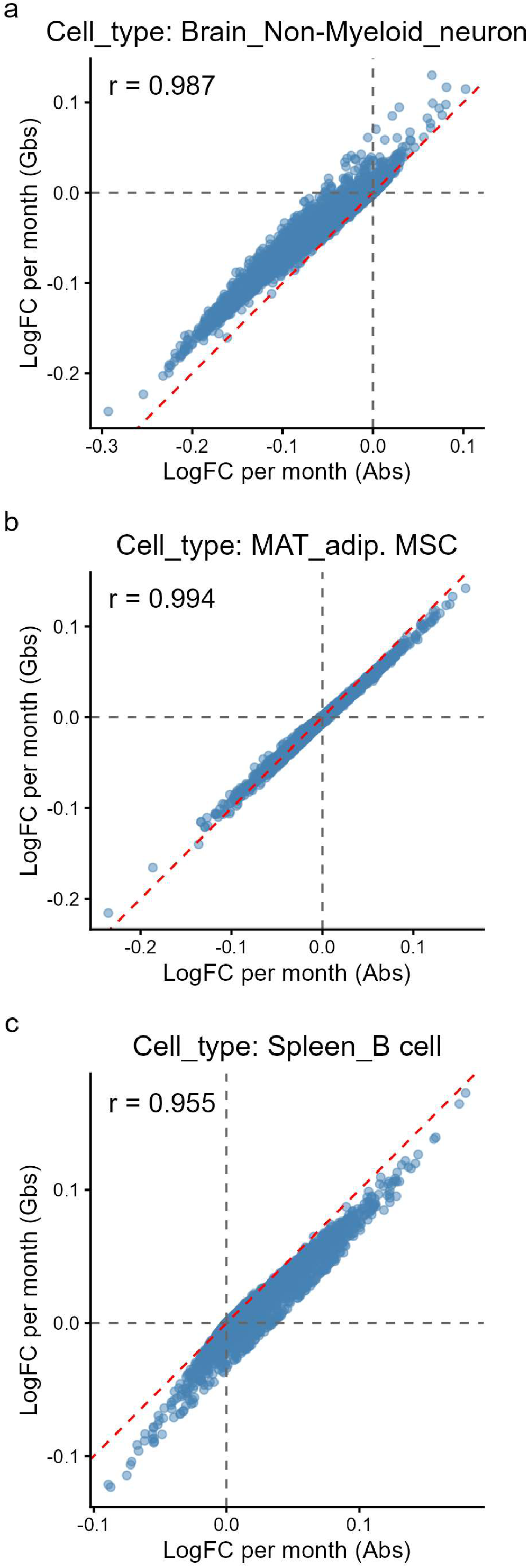
Comparison of absolute scaling with global scaling effect on logFC. **a)** Comparison of logFC with age estimated using absolute scaling and with global scaling in a cell type with reduced mRNA content (Brain_Non_mieloid_neuron). **b)** Comparison of logFC with age estimated using absolute scaling and with global scaling in a cell type with stable mRNA content (MAT_adipose_MSC). **c)** Comparison of logFC with age estimated using absolute scaling and with global scaling in a cell type with increased mRNA content (Spleen_B _cell).

**Figure S10.**
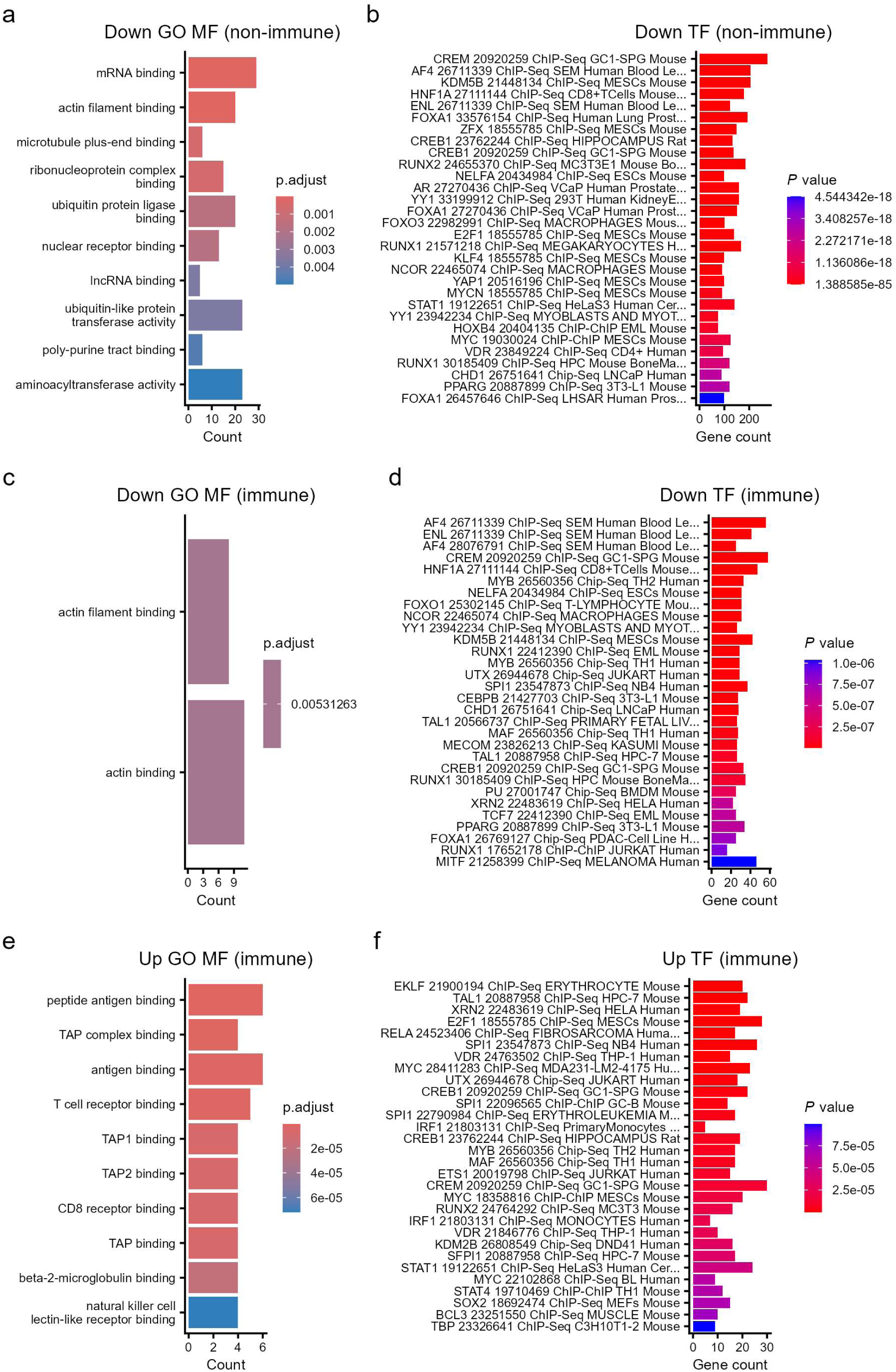
Functional enrichment of gene commonly changing with age. **a)** GO terms functional enrichment at molecular function in non-immune cells downregulated genes. **b)** Transcription factors enrichment analysis from enricher with ChEA_2022 regulons in non-immune cells downregulated genes. **c)** GO terms functional enrichment at molecular function in immune cells downregulated genes. **d)** Transcription factors enrichment analysis from enricher with ChEA_2022 regulons in immune cells downregulated genes. **e)** GO terms functional enrichment at molecular function in immune cells upregulated genes. **f)** Transcription factors enrichment analysis from enricher with ChEA_2022 regulons in immune cells upregulated genes.

**Figure S11.**
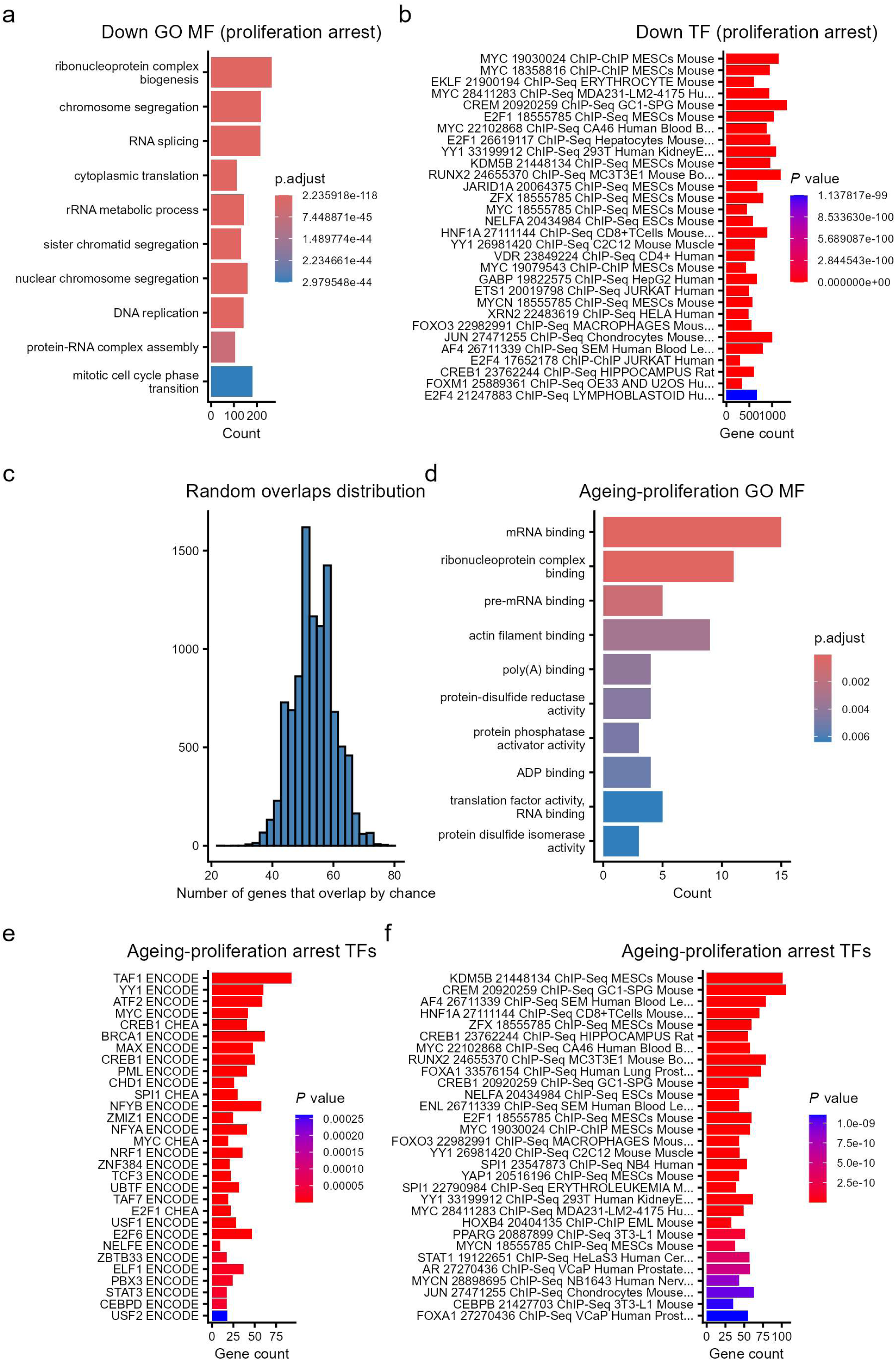
Proliferations arrest downregulated genes. **a)** GO terms functional enrichment at molecular function in proliferation arrest downregulated genes. **b)** Transcription factors enrichment analysis from enricher with ChEA_2022 regulons in proliferation arrest downregulated genes. **c)** distribution of the overlap random bootstrap (10 000, without replacement) of two sets of genes of the same size as the proliferating downregulated genes and age associated downregulated genes (p-value of 9.999e-05). **d)** GO terms functional enrichment at molecular function in the overlap genes that are downregulated in several cell types with age in non-immune cells with genes downregulated in the proliferation cessation. **e)** Transcription factors enrichment analysis from enricher with ENCODE and ChEA consensus regulons in the overlap genes that are downregulated in several cell types with age in non-immune cells with genes downregulated in the proliferation cessation. **f)** Transcription factors enrichment analysis from enricher with ChEA_2022 regulons in the overlap genes that are downregulated in several cell types with age in non-immune cells with genes downregulated in the proliferation cessation.

**Figure S12.**
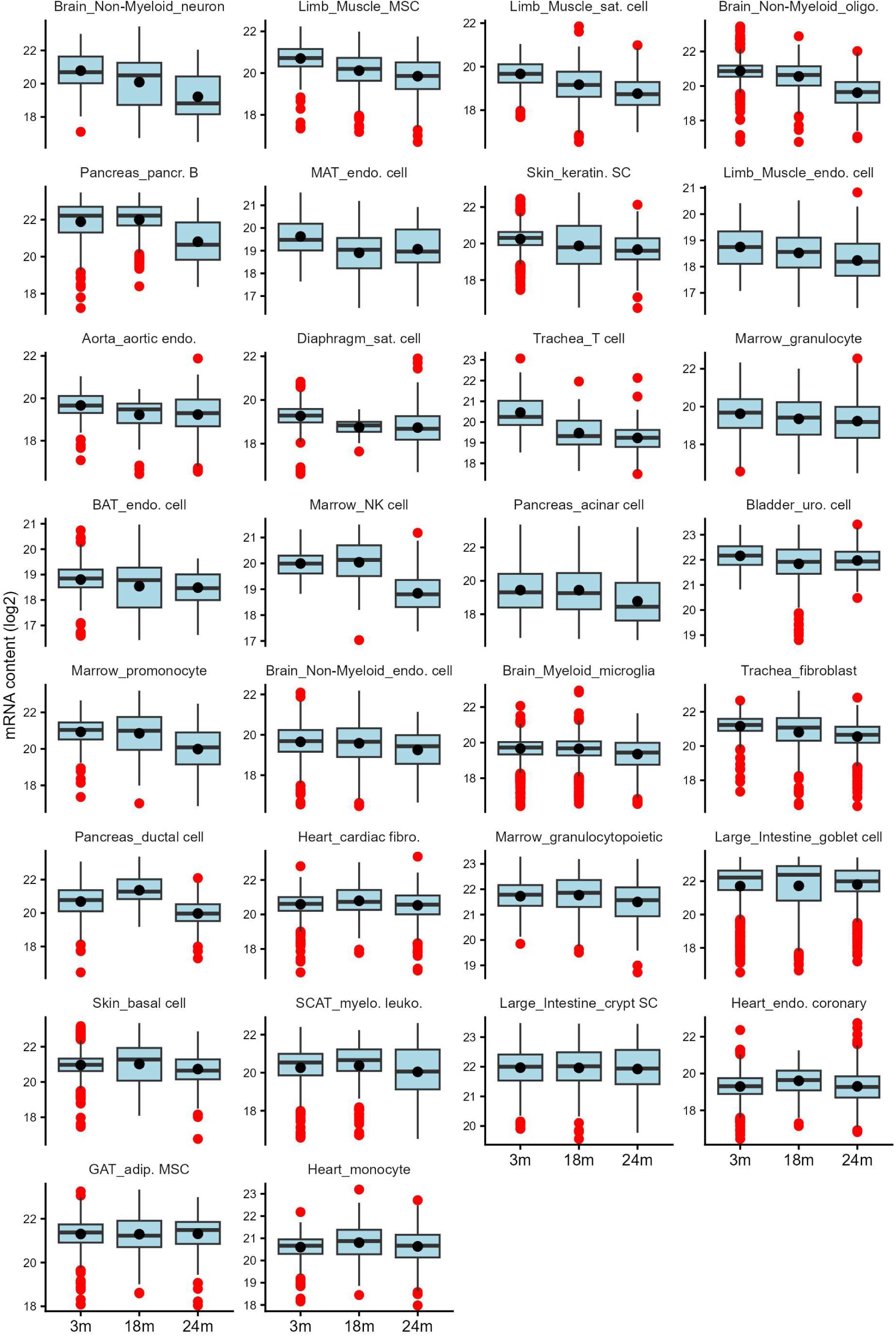
Boxplots of mRNA content by age. mRNA content by age in each cell type, ordered by mean mRNA content. First 30 cell types.

**Figure S13.**
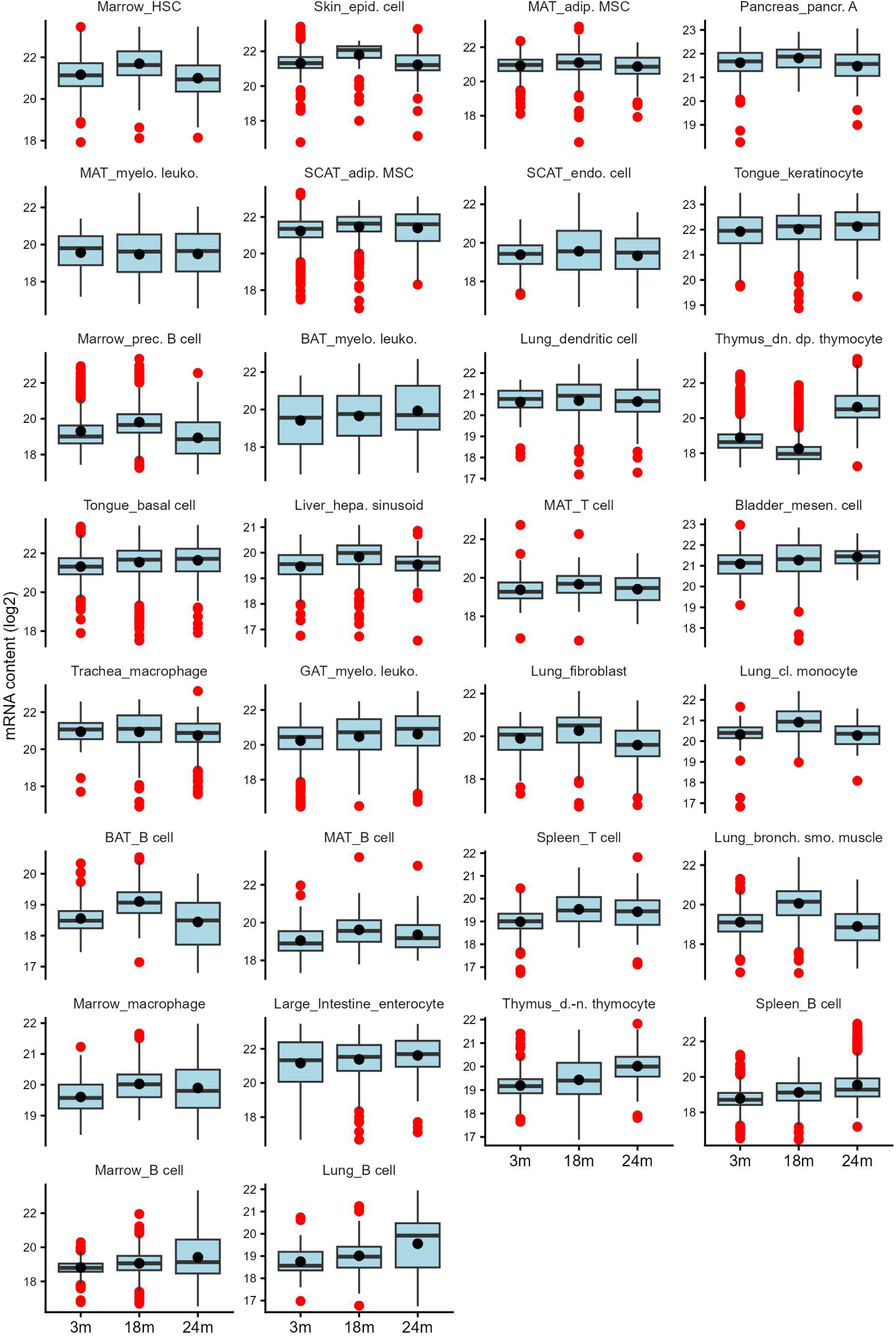
Boxplots of mRN content by age. mRNA content by age in each cell type, ordered by mean mRNA content. Last 30 cell types.

**Figure S14.**
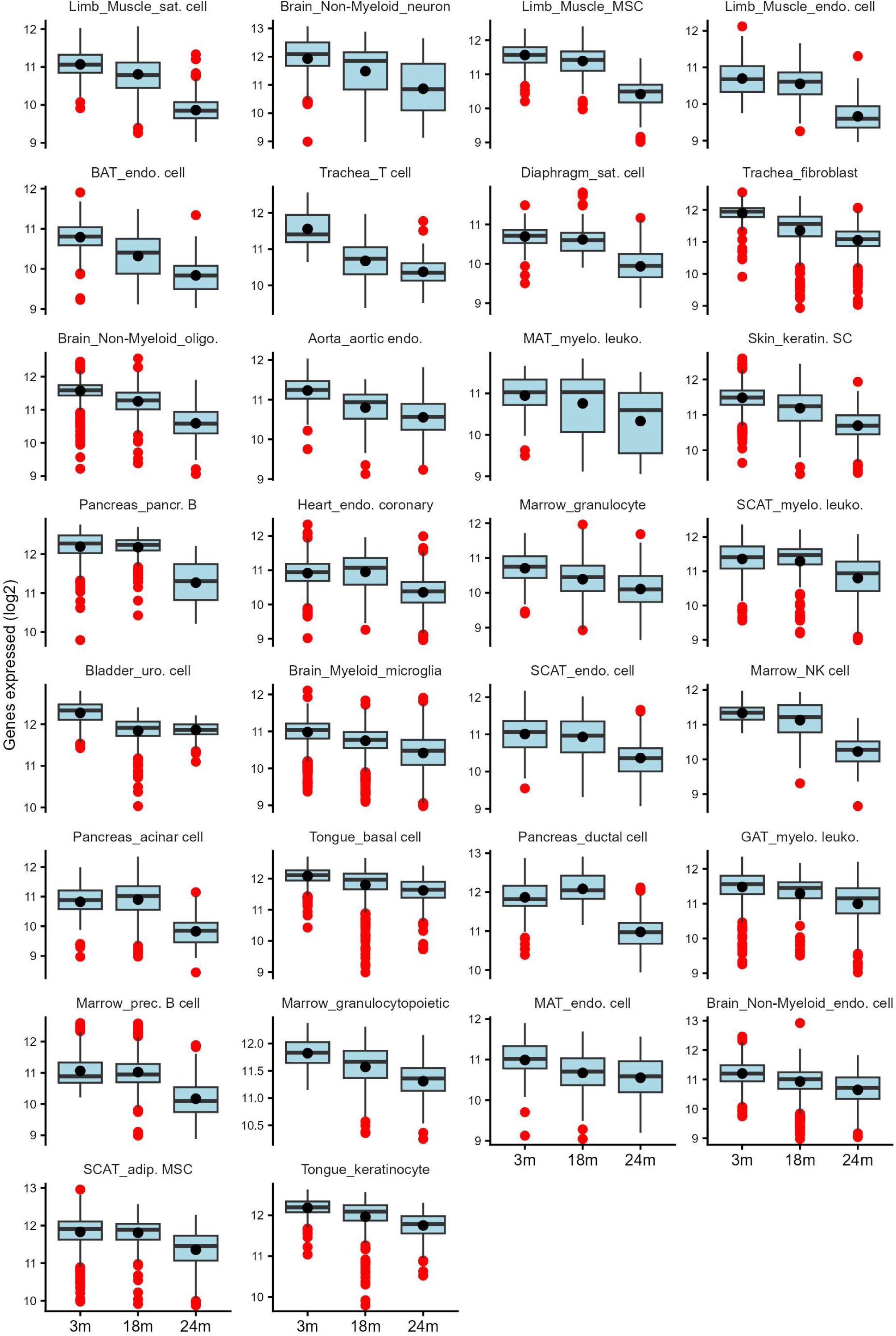
Boxplots of number of genes expressed by age. Number of genes by age in each cell type, ordered by mean number of genes expressed. First 30 cell types.

**Figure S15.**
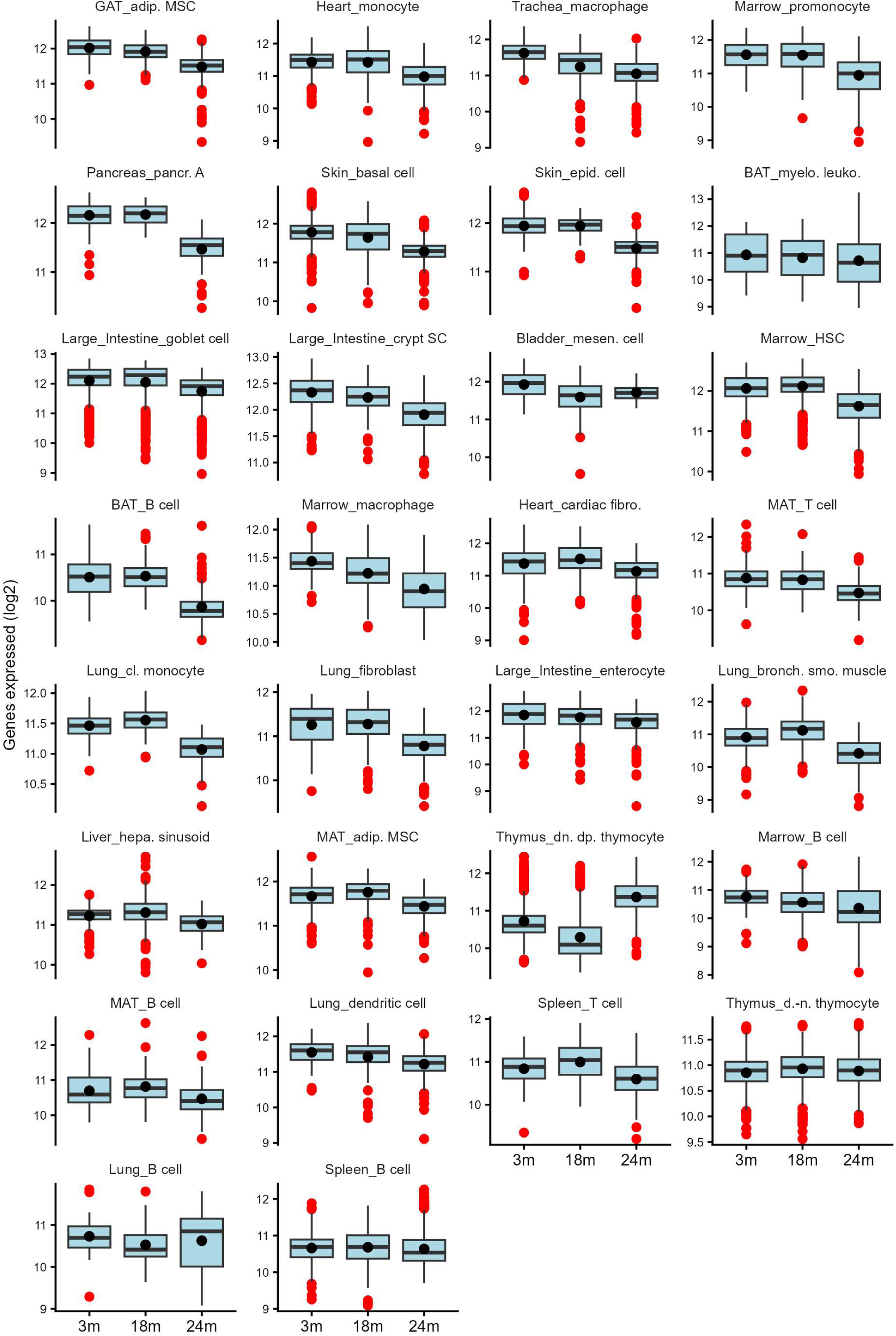
Boxplots of mRNA content by age. Number of genes by age in each cell type, ordered by mean number of genes expressed. Last 30 cell types.

## References

1. de Magalhães JP. 2024 Distinguishing between driver and passenger mechanisms of aging. Nat. Genet. 56, 204–211. (doi:10.1038/s41588-023-01627-0)

2. Osborne B et al. 2020 New methodologies in ageing research. Ageing Res. Rev. 62. (doi:10.1016/j.arr.2020.101094)

3. de Magalhães JP, Curado J, Church GM. 2009 Meta-analysis of age-related gene expression profiles identifies common signatures of aging. Bioinformatics 25, 875–881. (doi:10.1093/bioinformatics/btp073)

4. Huang Y, Zhu S, Yao S, Zhai H, Liu C, Han JDJ. 2025 Unraveling aging from transcriptomics. Trends in Genetics 41, 218–235. (doi:10.1016/j.tig.2024.09.006)

5. Quinn TP, Erb I, Gloor G, Notredame C, Richardson MF, Crowley TM. 2019 A field guide for the compositional analysis of any-omics data. Gigascience 8. (doi:10.1093/gigascience/giz107)

6. Almanzar N et al. 2020 A single-cell transcriptomic atlas characterizes ageing tissues in the mouse. Nature 583, 590–595. (doi:10.1038/s41586-020-2496-1)

7. Kim YK, Cho B, Cook DP, Trcka D, Wrana JL, Ramalho-Santos M. 2023 Absolute scaling of single-cell transcriptomes identifies pervasive hypertranscription in adult stem and progenitor cells. Cell Rep. 42. (doi:10.1016/j.celrep.2022.111978)

8. Berry S, Pelkmans L. 2022 Mechanisms of cellular mRNA transcript homeostasis. Trends Cell Biol. 32, 655–668. (doi:10.1016/j.tcb.2022.05.003)

9. Kuleshov M V. et al. 2016 Enrichr: a comprehensive gene set enrichment analysis web server 2016 update. Nucleic Acids Res. 44, W90–W97. (doi:10.1093/nar/gkw377)

10. Thomas MC, Chiang CM. 2006 The general transcription machinery and general cofactors. Crit. Rev. Biochem. Mol. Biol. 41, 105–178. (doi:10.1080/10409230600648736)

11. Panov KI, Friedrich JK, Russell J, Zomerdijk JCBM. 2006 UBF activates RNA polymerase I transcription by stimulating promoter escape. EMBO Journal 25, 3310–3322. (doi:10.1038/sj.emboj.7601221)

12. Sanij E et al. 2015 A novel role for the pol I transcription factor ubtf in maintaining genome stability through the regulation of highly transcribed pol II genes. Genome Res. 25, 201–212. (doi:10.1101/gr.176115.114)

13. Verheul TCJ, van Hijfte L, Perenthaler E, Barakat TS. 2020 The Why of YY1: Mechanisms of Transcriptional Regulation by Yin Yang 1. Front. Cell Dev. Biol. 8. (doi:10.3389/fcell.2020.592164)

14. Zhang MJ, Pisco AO, Darmanis S, Zou J. 2021 Mouse aging cell atlas analysis reveals global and cell type-specific aging signatures. Elife 10. (doi:10.7554/ELIFE.62293)

15. Jeffries AM, Yu T, Ziegenfuss JS, Tolles AK, Baer CE, Sotelo CB, Kim Y, Weng Z, Lodato MA. 2025 Single-cell transcriptomic and genomic changes in the ageing human brain. Nature 646, 657–666. (doi:10.1038/s41586-025-09435-8)

16. Kedlian VR et al. 2024 Human skeletal muscle aging atlas. *Nat*. Aging 4, 727–744. (doi:10.1038/s43587-024-00613-3)

17. Kim HS, Pickering AM. 2023 Protein translation paradox: Implications in translational regulation of aging. Front. Cell Dev. Biol. 11. (doi:10.3389/fcell.2023.1129281)

18. Bethlehem RAI et al. 2022 Brain charts for the human lifespan. Nature 604, 525–533. (doi:10.1038/s41586-022-04554-y)

19. Shaw SC, Dennison EM, Cooper C. 2017 Epidemiology of Sarcopenia: Determinants Throughout the Lifecourse. Calcif. Tissue Int. 101, 229–247. (doi:10.1007/s00223-017-0277-0)

20. Debès C et al. 2023 Ageing-associated changes in transcriptional elongation influence longevity. Nature 616, 814–821. (doi:10.1038/s41586-023-05922-y)

21. Tang W, Jørgensen ACS, Marguerat S, Thomas P, Shahrezaei V. 2023 Modelling capture efficiency of single-cell RNA-sequencing data improves inference of transcriptome-wide burst kinetics. Bioinformatics 39. (doi:10.1093/bioinformatics/btad395)

22. de Magalhães JP. 2025 An overview of contemporary theories of ageing. Nat. Cell Biol. 27, 1074–1082. (doi:10.1038/s41556-025-01698-7)

23. Milano L, Gautam A, Caldecott KW. 2024 DNA damage and transcription stress. Mol. Cell. 84, 70–79. (doi:10.1016/j.molcel.2023.11.014)

24. Gyenis A et al. 2023 Genome-wide RNA polymerase stalling shapes the transcriptome during aging. Nat. Genet. 55, 268–279. (doi:10.1038/s41588-022-01279-6)

25. Gulati GS et al. 2020 Single-cell transcriptional diversity is a hallmark of developmental potential. Science (1S7S). 367, 405–411. (doi:10.1126/science.aax0249)

26. Zatzman M et al. 2022 Widespread hypertranscription in aggressive human cancers. Sci. Adv. 8.

27. Chatsirisupachai K, Palmer D, Ferreira S, de Magalhães JP. 2019 A human tissue-specific transcriptomic analysis reveals a complex relationship between aging, cancer, and cellular senescence. Aging Cell 18. (doi:10.1111/acel.13041)

28. de Magalhães JP. 2025 The evolution of cancer and ageing: a history of constraint. Nat. Rev. Cancer 25, 873–880. (doi:10.1038/s41568-025-00861-4)

29. Wolf AM. 2021 The tumor suppression theory of aging. Mech. Ageing Dev. 200. (doi:10.1016/j.mad.2021.111583)

30. Lun ATL, Calero-Nieto FJ, Haim-Vilmovsky L, Göttgens B, Marioni JC. 2017 Assessing the reliability of spike-in normalization for analyses of single-cell RNA sequencing data. Genome Res. 27, 1795–1806. (doi:10.1101/gr.222877.117)

31. Robinson MD, Oshlack A. 2010 A scaling normalization method for differential expression analysis of RNA-seq data.

32. Grün D, Van Oudenaarden A. 2015 Design and Analysis of Single-Cell Sequencing Experiments. Cell 163, 799–810. (doi:10.1016/j.cell.2015.10.039)

33. Svensson V, Natarajan KN, Ly LH, Miragaia RJ, Labalette C, Macaulay IC, Cvejic A, Teichmann SA. 2017 Power analysis of single-cell rnA-sequencing experiments. Nat. Methods 14, 381–387. (doi:10.1038/nmeth.4220)

34. Lovén J, Orlando DA, Sigova AA, Lin CY, Rahl PB, Burge CB, Levens DL, Lee TI, Young RA. 2012 Revisiting global gene expression analysis. Cell. 151, 476–482. (doi:10.1016/j.cell.2012.10.012)

35. Raj A, van Oudenaarden A. 2008 Nature, Nurture, or Chance: Stochastic Gene Expression and Its Consequences. Cell. 135, 216–226. (doi:10.1016/j.cell.2008.09.050)

36. Jiang R, Sun T, Song D, Li JJ. 2022 Statistics or biology: the zero-inflation controversy about scRNA-seq data. Genome Biol. 23. (doi:10.1186/s13059-022-02601-5)

37. Squair JW et al. 2021 Confronting false discoveries in single-cell differential expression. Nat. Commun. 12. (doi:10.1038/s41467-021-25960-2)

38. Zhu D, Arnold M, Samuelson BA, Wu JZ, Mueller A, Sinclair DA, Kane AE. 2024 Sex dimorphism and tissue specificity of gene expression changes in aging mice. Biol. Sex Differ. 15. (doi:10.1186/s13293-024-00666-4)

39. Luciano A, Robinson L, Garland G, Lyons B, Korstanje R, Di Francesco A, Churchill GA. 2024 Longitudinal fragility phenotyping contributes to the prediction of lifespan and age-associated morbidity in C57BL/6 and Diversity Outbred mice. Geroscience 46, 4937–4954. (doi:10.1007/s11357-024-01226-9)

40. Amezquita RA et al. 2020 Orchestrating single-cell analysis with Bioconductor. Nat. Methods 17, 137–145. (doi:10.1038/s41592-019-0654-x)

41. Durinck S, Moreau Y, Kasprzyk A, Davis S, De Moor B, Brazma A, Huber W. 2005 BioMart and Bioconductor: A powerful link between biological databases and microarray data analysis. Bioinformatics 21, 3439–3440. (doi:10.1093/bioinformatics/bti525)

42. Andrews N et al. 2021 An unsupervised method for physical cell interaction profiling of complex tissues. Nat. Methods 18, 912–920. (doi:10.1038/s41592-021-01196-2)

43. Montserrat-Ayuso T, Esteve-Codina A. 2024 High content of nuclei-free low-quality cells in reference single-cell atlases: a call for more stringent quality control using nuclear fraction. BMC Genomics 25. (doi:10.1186/s12864-024-11015-5)

44. Lagger C, Ursu E, Equey A, Avelar RA, Pisco AO, Tacutu R, de Magalhães JP. 2023 scDiffCom: a tool for differential analysis of cell–cell interactions provides a mouse atlas of aging changes in intercellular communication. *Nat*. Aging 3, 1446–1461. (doi:10.1038/s43587-023-00514-x)

45. Germain PL, Lun A, Macnair W, Robinson MD. 2021 Doublet identification in single-cell sequencing data using scDblFinder. F1000Res. 10, 979. (doi:10.12688/f1000research.73600.1)

46. Scialdone A, Natarajan KN, Saraiva LR, Proserpio V, Teichmann SA, Stegle O, Marioni JC, Buettner F. 2015 Computational assignment of cell-cycle stage from single-cell transcriptome data. Methods 85, 54–61. (doi:10.1016/j.ymeth.2015.06.021)

47. Locard-Paulet M, Palasca O, Jensen LJ. 2022 Identifying the genes impacted by cell proliferation in proteomics and transcriptomics studies. PLoS Comput. Biol. 18. (doi:10.1371/journal.pcbi.1010604)

48. Aibar S et al. 2017 SCENIC: Single-cell regulatory network inference and clustering. Nat. Methods 14, 1083–1086. (doi:10.1038/nmeth.4463)

49. Wu T et al. 2021 clusterProfiler 4.0: A universal enrichment tool for interpreting omics data. Innovation 2. (doi:10.1016/j.xinn.2021.100141)

50. Kuleshov M V. et al. 2016 Enrichr: a comprehensive gene set enrichment analysis web server 2016 update. Nucleic Acids Res. 44, W90–W97. (doi:10.1093/nar/gkw377)

